# A rapid transfer of virions coated with heparan sulfate from the ECM to CD151 defines an early step in the human papillomavirus infection cascade

**DOI:** 10.1101/2025.04.17.649398

**Authors:** Annika Massenberg, Yahya Homsi, Carl Niklas Schneider, Snježana Mikuličić, Tatjana Döring, Luise Florin, Thorsten Lang

## Abstract

Human Papillomaviruses (HPVs) are the underlying cause of several types of cancer, albeit they are mostly known for their association with cervical carcinoma. The virions reach their target cells through a break in the epithelial barrier. After binding to heparan sulfate (HS) of the extracellular matrix (ECM), they are recruited via actin-dependent mechanisms to the cell surface where they co-internalize with the entry factor CD151.

The *in vivo* occurring active recruitment from the ECM to the cell surface may be bypassed in cell culture where virions reach the cell surface simply by passive diffusion. To specifically investigate these early events of the infection cascade, we use HaCaT keratinocytes as they produce a robust ECM enabling for abundant virion binding to ECM components such as HS before transfer to cell surface receptors and infection. Employing microscopy, we focus on the basal membrane that for virions is difficult to access by diffusion. We block the active recruitment from ECM attachment sites to the cell body, release the blocking, and monitor the association of virions with CD151 or HS. We observe quick virion recruitment from the ECM to the cell body within 15 min. During recruitment, virions associate with the tetraspanin CD151 present at the cell border or at filopodia. These virions are decorated with HS, which they lose in the next few hours, presumably prior to endocytosis.

Our observations reveal a rapid step in the HPV infection cascade: the transfer of HS-coated virions from the ECM to CD151. This step is too fast to account for the asynchronous uptake of HPVs which is likely driven by glycan- and capsid processing.

## Introduction

Already in the 80s, Harald zur Hausen proposed a role of human papillomaviruses in cancer (***zur Hausen, 2009***). Since then, five more classes of oncogenic viruses have been identified (***Galati et al., 2024a***). To date, it is assumed that more than 10% of the worldwide human cancer burden is associated with infectious agents (***Galati et al., 2024a***), from which about a half is caused by *Papillomaviridae* (***Martel et al., 2017***). Thus, the understanding of viral entry strategies has implications going beyond the classical treatment of acute viral infections.

Human papillomaviruses are small, non-enveloped viruses with a diameter of ≈55 nm. The icosahedral capsid is mainly composed of pentameric L1 capsomers. Together with fewer L2 capsid proteins, capsomers surround a histone core bearing a circular double-stranded DNA (***Baker et al., 1991; Ozbun and Campos, 2021***). From the more than 200 phylogenetically classified HPV genotypes, the most oncogenic ones are HPV16 and HPV18 (***Galati et al., 2024b***), which are responsible for about 70 - 80% of the cervical cancer cases (***Christiansen et al., 2015***). In addition, they cause other severe cancers such as anogenital, head and neck tumors (***Doorbar et al., 2012***).

For papillomavirus infection, a disruption of the epithelial barrier is a prerequisite, through which virions reach mitotically active basal cells of the epithelia (***Ozbun, 2019***). Here, virions bind to the linear polysaccharide heparan sulfate (HS) that is present in the extracellular matrix (ECM) and at the plasma membrane surface. HS is covalently linked to proteins forming so called heparan sulfate proteoglycans (HSPGs). Positively charged and polar amino acid residues of the L1 capsid protein form multiple heparan sulfate binding sites that interact with negative charges of HS, resulting in a strong bond (***Dasgupta et al., 2011; Giroglou et al., 2001; Joyce et al., 1999; Knappe et al., 2007; Surviladze et al., 2015***). While in cell culture virions bind to HS on both the cell surface and the ECM, it has been suggested that *in vivo* they bind predominantly to HS of the extracellular basement membrane (***Day and Schelhaas, 2014; Kines et al., 2009; Ozbun and Campos, 2021; Schiller et al., 2010***). In any case, the link between the linear polysaccharide and virions must be disrupted before they can bind to a yet unknown secondary receptor on the cell surface, followed by internalization (***Ozbun and Campos, 2021***).

The strong electrostatic bonding precludes dissociation as a virion release mechanism. Two alternatives are discussed, that mutually are not exclusive. In the so-called priming model, binding of HS to the capsid results in capsid enlargement and softening (Feng et al., 2024), followed by the exposure and cleavage of L1 by kallikrein-8 (KLK8) (***Cerqueira et al., 2015***), exposure of L2 by cyclophilin (***Bienkowska-Haba et al., 2009***), and L2 cleavage by furin (***Richards et al., 2006***). After these structural modifications of the capsid surface the so-called ‘primed’ virion is able to bind to the secondary receptor. In an alternative model, the HS-virion bond persists. However, heparanases and proteinases cleave HS/HSPGs into fragments. As a result, albeit still bound to the virion surface, the now fragmented HS/HSPG no longer anchors the virion to the ECM (***Surviladze et al., 2012; Surviladze et al., 2015***). Next, released virions may reach cell surface receptors simply by passive diffusion. However, active recruitment is possible as well. For instance, PsVs migrate along actin-rich protrusions from the ECM towards the cell body (***Schelhaas et al., 2008; Smith et al., 2008***). Moreover, cells may reach virions by migrating onto an ECM lawn decorated with virions.

While it is generally agreed on that HS is a primary virion attachment site, the molecular identity of the secondary receptor on the main cell body surface is unknown. The secondary receptor complex is most likely of multimeric nature rather than a single molecular component. Possible candidates are proteins known to be crucial for cell entry, as the tetraspanin CD151 (***Mikuličić et al., 2019; Scheffer et al., 2013; Spoden et al., 2008***), integrin-α6 (Itgα6) (***Evander et al., 1997; Yoon et al., 2001***), growth factor receptors (***Mikuličić et al., 2019; Surviladze et al., 2012***), and the annexin A2 heterotetramer (***Dziduszko and Ozbun, 2013; Woodham et al., 2012***). From these molecules, the tetraspanin CD151 could play a coordinating role, as tetraspanins organize viral entry platforms in many types of viral infections, including infections with coronavirus, cytomegalovirus, hepatitis C virus, human immunodeficiency virus, human papilloma virus, and influenza virus (***Bruening et al., 2018; Earnest et al., 2015; Florin and Lang, 2018; Hantak et al., 2019; Hochdorfer et al., 2016; Scheffer et al., 2013; Zona et al., 2013***). Such tetraspanin entry platforms could form in a slow and stochastic fashion, which would provide an explanation for the asynchronous virion uptake with half-times of above 10 h (***Becker et al., 2018***). However, it is unclear whether virions associate with CD151 already in the moment of virion transfer from the ECM to the cell surface, or in a subsequent step, e.g. in preparation of endocytosis.

Here, we explore the question of a possible link between active virion recruitment to the cell surface and CD151 association. We employ the cell permeable mycotoxin and actin polymerization inhibitor cytochalasin D (CytD). Using a human keratinocyte cell line (HaCaT cells), we find that CytD preserves HPV16 PsVs in the ECM, noticed as PsV accumulations adjacent to the cell periphery. This blocking of the virus transfer is accompanied by co-accumulation of HS in the ECM area. Upon CytD removal, HS-decorated PsVs get from the ECM to the cell body where they associate with CD151. The association of PsVs with CD151 persists within the next few hours, whereas the HS coat is stripped off, and CD151 is observed to agglomerate. These findings distinguish an early step in the infection cascade, the association with CD151 in the moment the virion establishes contact to the cell surface.

## Results

### Cytochalasin D arrests the active recruitment of HPV16 PsVs from the extracellular matrix to the basal cell membrane

The molecular surface of PsVs is immunologically indistinguishable from HPV virus particles (***Ozbun and Campos, 2021***), which makes them a widely used tool for studying host cell entry. In this study, we employ HPV16 pseudovirions with an encapsidated luciferase reporter plasmid under the control of the HPV16 promotor (***Wüstenhagen et al., 2018***). Hence, instead of viral DNA a luciferase encoding plasmid enters the cell, enabling the analysis of the infection rate via the luciferase activity. Moreover, the plasmids are composed of nucleotides to which fluorophores can be coupled by click-chemistry. This allows for their microscopic detection by fluorescence microscopy, alternatively to immunolabeling of the L1 capsid protein.

PsVs that bind to the ECM at sites distal from the cell body are unable to establish direct contact with entry receptors, until the cell migrates onto them or they are transported along cell protrusions towards the cell body (***Schelhaas et al., 2008; Smith et al., 2008***). Both cell migration and protrusion transport depend on actin dynamics (***Schaks et al., 2019***). We aimed for blocking these active recruitment mechanisms in HaCaT cells, a cell line that is widely used as a cell culture model for HPV infection. They resemble primary keratinocytes in several key aspects: they are not virally transformed and produce large amounts of ECM, promoting interactions between viruses and ECM components and thereby facilitating infection (***Bienkowska-Haba et al., 2018; Gilson et al., 2020***). In addition, subconfluent HaCaT cells form filopodia and filopodial transport is used for the recruitment of ECM-bound virus particles to the cell body *(**Schelhaas et al., 2008, Smith et al., 2008**)*. Together, these features make HaCaT cells a suitable model for studying active PsV recruitment from the ECM to the cell surface.

We incubate cells for 5 h with PsVs in the absence or presence of 10 µg/ml (19.7 µM) CytD, a concentration that stops cell migration after a few minutes (***Peng et al., 2011***; using 10 µM CytD). Moreover, in HeLa cells, retrograde transport of virions is sensitive to CytD (2 µM) (***Schelhaas et al., 2008***). Hence, under CytD, active PsV recruitment should be largely inhibited. Frequently, we observe patches of confluent cells which are common to HaCaT cells. Cells at the center of these patches are dismissed during imaging, because hardly any PsVs are bound to their basal membrane, indicating that PsVs do rather not reach this area by passive diffusion. Instead, we focus on isolated HaCaT cells or cells at the periphery of cell patches. At these cells, we find more PsVs per cell than one would expect from the employed ≍ 50 viral genome equivalents (vge) per cell, indicating that PsVs are unequally distributed between the cells. Moreover, in particular after CytD treatment, these PsVs usually are not homogenously distributed around the cell but rather concentrate at one region (Figures 1 to 4). In later experiments (Figures 5 to 8), we investigate the recruitment of PsVs from these regions, defining ROIs for analysis that cover PsVs at the periphery and the cell body (see Supplementary Figures 5A and 8A).

**Figure 1.**
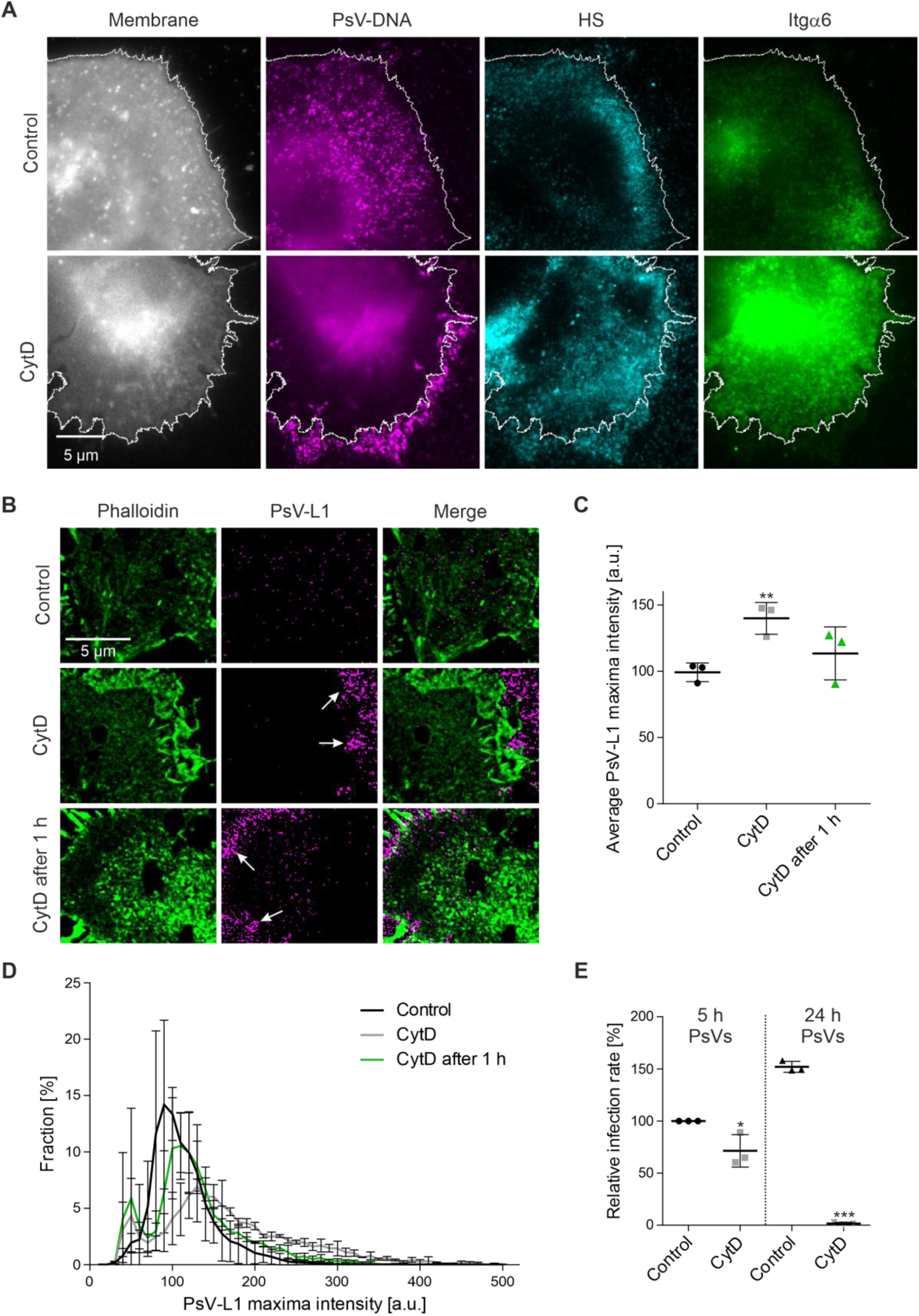
CytD arrests PsV recruitment from the ECM to the cell body. (A) In the absence (Control) or presence of 10 µg/ml CytD (CytD), HaCaT cells were incubated with PsVs at 37 °C for 5 h. Then, cells were fixed, washed and stained with the cell membrane dye TMA-DPH (gray lookup table (LUT)). PsVs (magenta LUT) were visualized through coupling a dye (6-FAM Azide) to the encapsidated plasmid by click-chemistry. Indirect immunolabeling was employed for staining of HS (AlexaFluor 594; cyan LUT) and Itgα6 (STARRED; green LUT). Imaging was realized with epi-fluorescence microscopy. White lines delineate the main cell body; lines were created with reference to the TMA-DPH membrane staining. (B) Same pre-treatment of cells as in (A), with an additional condition where CytD was added 1 h after the PsVs (CytD after 1 h). Prior to fixation, membrane sheets were generated and F-actin was stained with phalloidin coupled to iFluor488 (green LUT). The capsid protein L1 of the PsVs and CD151 were stained by immunofluorescence using primary antibodies in combination with AlexaFluor 594-labelled (L1, magenta LUT) and STAR RED-labelled (CD151, not shown in this figure for clarity reasons) secondary antibodies. Images of phalloidin and L1 staining were acquired in the confocal and STED mode of a STED microscope, respectively. Arrows in the PsV-L1 images point towards accumulated PsVs that after CytD are more frequently observed than in the control (see text). (C) Analysis of images as shown in (B) using ROIs covering the whole image. PsV maxima were detected and their intensities were quantified in a circular 125 nm diameter region of interest (ROI), followed by background correction. Values are given as means ± SD (n = 3; one biological replicate includes per condition the average of 14 - 15 analyzed membranes (intensity values of one membrane sheet were averaged) with altogether at least 1000 maxima intensity values). (D) PsV maxima intensity distribution of the data in (C). The fraction of PsVs, expressed in percent, is plotted as histogram (10 a.u. bins) against the maxima intensity. Values are given as means ± SD (n = 3). (E) HaCaT cells were treated either for 5 h or 24 h with PsVs, without (Control) or with 10 µg/ml CytD (CytD). In case of the 5 h incubation, after removal of the PsVs/CytD cells were incubated for another 19 h in medium (in total 24 h). After a total of 24 h incubation, the luciferase activity of lysed cells was measured, yielding the infection rate that was normalized to LDH, resulting in the normalized infection rate. The normalized infection rate was further related to the mean normalized infection rate of the 5 h control, set to 100%, yielding the relative infection rate. Values are given as means ± SD (n = 3 biological replicates; the value of one biological replicate is the average of three technical replicates). (C) and (E). Statistical differences between Control and CytD was analyzed by using the two-tailed, unpaired student’s *t*-test (n = 3, for details see methods). a.u., arbitrary units.

In Figure 1, after the 5 h incubation, cells were fixed and stained by a membrane marker, the dye TMA-DPH, to mark the main cell body (Figure 1A, gray lookup table (LUT)). Additionally, PsVs were visualized by click-chemistry (magenta LUT), and HS (cyan LUT) and the membrane protein Itgα6 (green LUT) were stained by antibodies. In order to monitor only cell surface events, we stain the cells without prior cell permeabilization (still, fixation perforates the cell membrane to some extent). In the control, PsVs locate strongly in the cell body area (Figure 1A, upper row; see white outline that is based on TMA-DPH staining). In contrast, when CytD was present, the cell body area is largely devoid of PsVs (Figure 1A, lower row; please note that the large central bright area is mainly caused by autofluorescence). Instead, PsVs often accumulate adjacent to the cell body. This adjacent region, that is up to several µm wide and rich in HS, likely includes the ECM. We conclude that upon inhibition of actin-dynamics PsVs do not efficiently reach the basal cell membrane but rather remain accumulated in the ECM area. In contrast, in the control they are actively recruited to the basal cell membrane. Please note that under all conditions PsVs will bind to receptors located outside of the imaged basal membrane, e.g., the entire upper cell membrane, but these PsVs are not visible in the micrographs and therefore not included in the analysis.

Comparing in Figure 1A the PsV brightness in the upper (control, magenta LUT) and lower (CytD, magenta LUT) panel suggests that the PsV amounts do not differ greatly. This is surprising, as CytD is not reported to inhibit enzymes involved in capsid modification or HS/HSPGs processing, after which the PsVs supposedly leave the ECM. Provided PsVs processed under CytD would leave the ECM, no major difference in the number of PsVs at the periphery between cells that were treated with or without CytD for 5 h is expected. Therefore, it appears that primed PsVs remain ECM associated. Due to interference from autofluorescence (Figure 1A, large magenta areas in the centers), a quantification of the PsVs with and without CytD is not possible. To study PsVs in the absence of autofluorescence, we employed unroofed cells (***Heuser, 2000***), also referred to as membrane sheets. After pre-treatment of cells as above, membrane sheets are generated by brief ultrasound pulses that remove the upper cellular parts (and thus any intracellular autofluorescence), leaving behind the extracellular matrix and the basal membrane along with the bound PsVs. During imaging, membrane sheets are identified via staining of F-actin with fluorescently labelled phalloidin (Figure 1B, green LUT) that marks the main cell body cortex together with lamellipodia and filopodia. In this experiment we again observe that PsVs, not always but more frequently, after CytD treatment accumulate adjacent to the isolated basal membrane (Figure 1B, magenta LUT, see arrows) that is defined by F-actin staining (Figure 1B, green LUT).

The PsV maxima density (Figure 1B, magenta LUT, detected as local maxima of antibody stained L1) and the PsV intensity are quantified in an area covering the cell body and the periphery. Compared to the control, CytD reduces the PsV maxima density by 26% (the PsV maxima density in the control and CytD treated cells is 0.7 and 0.52 PsVs/µm^2^, respectively), whereas the maxima intensity increases by 41% (Figure 1C). A histogram of the PsV maxima intensities illustrates that CytD broadens the intensity distribution towards brighter PsVs (Figure 1D, compare black and gray trace). The increase of the PsV intensity after CytD treatment can also be appreciated comparing the upper and middle/lower L1 images in Figure 1B (shown at the same settings of brightness and contrast). As CytD is unlikely to act directly on PsVs (e.g., by fusing them to larger and brighter particles), we assume that the underlying cause of brighter maxima is the resolution limit of the microscope. The densely accumulated PsVs are no longer resolved, overlap, and by this merge to brighter spots. These poorly resolved maxima are on average 41% brighter but of them there are 26% fewer. The decrease in maxima density is in the order of magnitude of the increase in intensity, which yields a roughly similar total signal in both conditions. Hence, active recruitment apparently thins out the PsV accumulations by translocating PsVs from the ECM to the cell body.

We addressed whether PsVs would still be accumulated when adding CytD 1 h after the PsVs, rather than adding them simultaneously (Figure 1B, lower panel, CytD after 1 h). As in the CytD condition, accumulated PsVs adjacent to the basal membrane are present more frequently than in the control (Figure 1B, arrows). The PsV maxima intensity distribution (Fig. 1D, green trace) is slightly broader when compared to the control, but narrower than with CytD present for the entire time (Figure 1D), suggesting that PsV binding and recruitment take more than 1 h.

Figure 1 indicates that under CytD, when PsVs are not actively collected from the cells, they remain trapped in the ECM next to the cell body. However, for several reasons it cannot be determined exactly how efficient the preservation of accumulated PsVs is. First, our assay is not precise for technical reasons, as the above-mentioned limited resolution results in an underestimation of the number of PsVs the stronger they are accumulated. Second, we do not know how many PsVs in the control are endocytosed during the 5 h of incubation, which is an important number, as adding theses PsVs to the ones being present would be the adequate reference value for comparison to CytD. Moreover, it should be noted that both the degree of PsV accumulation as well as the number of PsVs are highly variable. Even in the control, we occasionally find cells with accumulated PsVs at the periphery, while after CytD we not always encounter accumulated PsVs. This causes a large variability within the data. Still, we consider larger PsV accumulations as a typical effect of CytD incubation (please note that for illustration of this effect we show in the Figures such examples). In any case, we conclude that within the 5 h incubation period with CytD a large fraction of ECM associated PsVs is not able to reach the cell body as in the control, but remains located in its periphery.

### The blocking of PsV translocation by cytochalasin D is reversible

CytD not only arrests the recruitment of PsVs to the cell body but is known to block other actin-dependent processes, which strongly affects the physiology of the cell. Therefore, we investigated whether PsVs would proceed normally on their infection pathway once CytD is washed off. In order to allow for recovery from CytD, cells were washed after the 5 h PsV/CytD treatment, then incubated for another 19 h and the infection rate was determined by measuring the luciferase activity in the cell lysate. CytD reduced luciferase activity by 29%, as compared to control cells incubated only with PsVs (Figure 1E; left). When PsVs were not washed off after 5 h but left for the entire 24 h incubation, the infection rate increased by 52%, whereas continuous treatment with CytD blocks infection (Figure 1E; right). The latter is in line with previous studies (***Schelhaas et al., 2008; Selinka et al., 2002; Selinka et al., 2007; Spoden et al., 2013***) showing that CytD is a strong inhibitor of HPV infection. In any case, PsVs apparently are able to proceed on the infection pathway upon removal of CytD. The reduced infection rate of 29% can be explained by the 5 h blocking at the beginning of the incubation period that should delay the time course of infection. Altogether, we propose that CytD is suitable for transiently arresting PsVs in a state between primary attachment to HS and cell body receptor binding.

Next, we investigated the onset of active PsV recruitment after CytD wash off. Cells were treated with PsVs and CytD for 5 h, washed, and incubated further without PsVs/CytD for 0 min, 30 min or 60 min, followed by fixation and staining for PsVs (Figure 2A, magenta LUT) and F-actin (Figure 2A, green LUT). The antibody staining against PsVs, like above the click-chemistry staining, is again highly variable. As a measure of recruitment to the cell body, we calculated the Pearson correlation coefficient (PCC) between PsV-L1 and F-actin staining. The PCC quantifies the similarity between two variables, in this case the pixel values of the two images. The PCC results in the value of 1 if images are identical and -1 for an image and its negative. After 5 h of CytD, PsVs often are found accumulated, as seen before, at the edge of the F-actin staining (Figure 2A). The distal PsVs and the F-actin staining partially exclude each other, thus at 0 min we obtain a negative PCC (Figure 2A and D). After 30 min and 60 min, PsVs mostly overlap with F-actin stained areas, which results in positive PCCs (Figure 2A and D). The time course of the change in PCC values suggests an onset of recruitment after CytD removal within 30 min.

**Figure 2.**
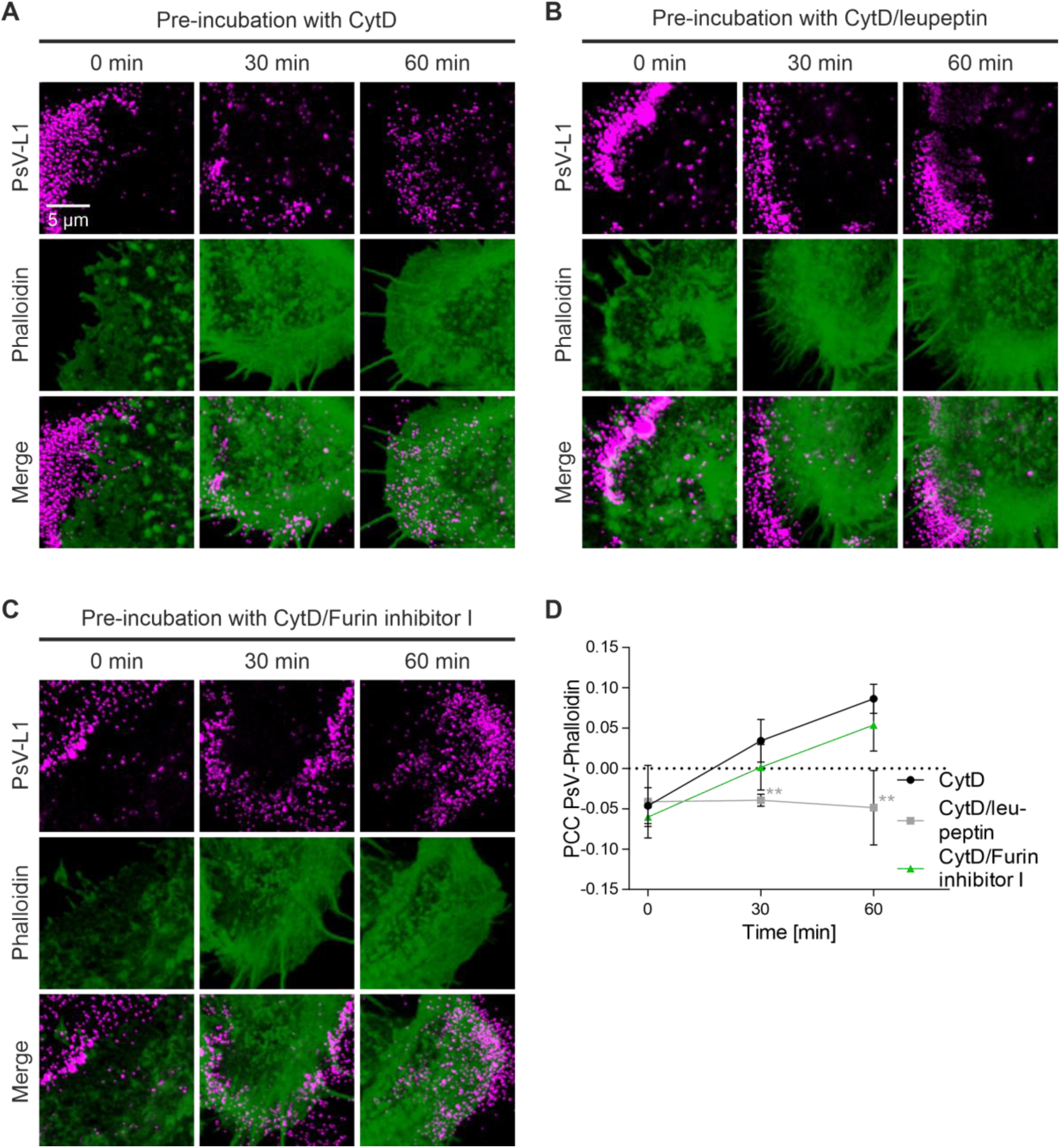
Recruitment of PsVs from the ECM to the cell body requires PsV priming. HaCaT cells were pre-incubated for 5 h at 37 °C with PsVs, in the presence of (A) 10 µg/ml CytD (CytD), (B) 10 µg/ml CytD and 100 µM leupeptin (CytD/leupeptin), or (C) 10 µg/ml CytD and 5 µM Furin inhibitor I (CytD/Furin inhibitor I). Afterwards, cells were washed and incubated without PsVs/inhibitors further for 0 min, 30 min or 60 min, before they were fixed and stained by indirect immunofluorescence for L1 (STAR GREEN, magenta LUT) and for F-actin by iFluor647-labelled phalloidin (green LUT). PsVs-L1 and F-actin staining were imaged in the confocal mode of a STED microscope. (D) For determination of the Pearson correlation coefficient (PCC) between PsV-L1 (magenta LUT) and Phalloidin (green LUT), we placed large ROIs onto the images that covered mainly the cell body but included parts of the cell periphery as well. The PCC was plotted over time. Values are given as means ± SD (n = 3 biological replicates). Statistical difference between CytD and CytD/inhibitors was analyzed by using the two-tailed, unpaired student’s *t*-test (n = 3, for details see methods).

### Confirming in our assay previously proposed steps in HPV infection

As outlined above, PsVs bind tightly to their primary attachment site until the capsid surface undergoes structural changes, which involves modifications of L1 and L2. Consequently, we expect that inhibition of L1 processing during the CytD incubation should inhibit PsV recruitment after CytD removal (Figure 2A and D). To test for this possibility, as employed in earlier studies, the protease inhibitor leupeptin was used to inhibit proteases including KLK8 which is required for L1 cleavage (***Cerqueira et al., 2015***). Employing this inhibitor, the PCC between PsV-L1 and F-actin staining remains negative after CytD removal, showing that for active recruitment indeed the prior action of proteases is a prerequisite (Figure 2B and D). Moreover, the experiment suggests that without PsV priming the PCC between PsV-L1 and F-actin does not increase, for instance, due to cell spreading after CytD removal. In contrast, inhibition of L2 cleavage by a furin specific inhibitor has no effect on the PCC (Figure 2C and D). However, it should be noted that we occasionally observe PsVs not undergoing complete translocation. Instead, they remain accumulated at the border of the F-actin stained area (for example see Figure 2C, 60 min). This results in an increase of the PCC as in complete translocation, explaining why the PCC changes like in the control, despite of a furin effect. Hence, furin may have some effect on a later recruitment step that, however, is undetected in this type of analysis.

As outlined above, during the 5 h incubation with CytD, proteases in the ECM are expected to cleave HS chains. These cleavage products should be able to diffuse out of the ECM, unless they remain associated with non-translocating PsVs, present in particular under CytD. Using an antibody that reacts with an epitope present in native heparan sulfate chains, we find that only after CytD and if PsVs were present, the level of HS staining is significantly increased (Figure 3B). As shown in Figure 3A, this stronger HS staining is observed in areas with PsVs (open arrows) and as well in PsV free areas (closed arrows).

**Figure 3.**
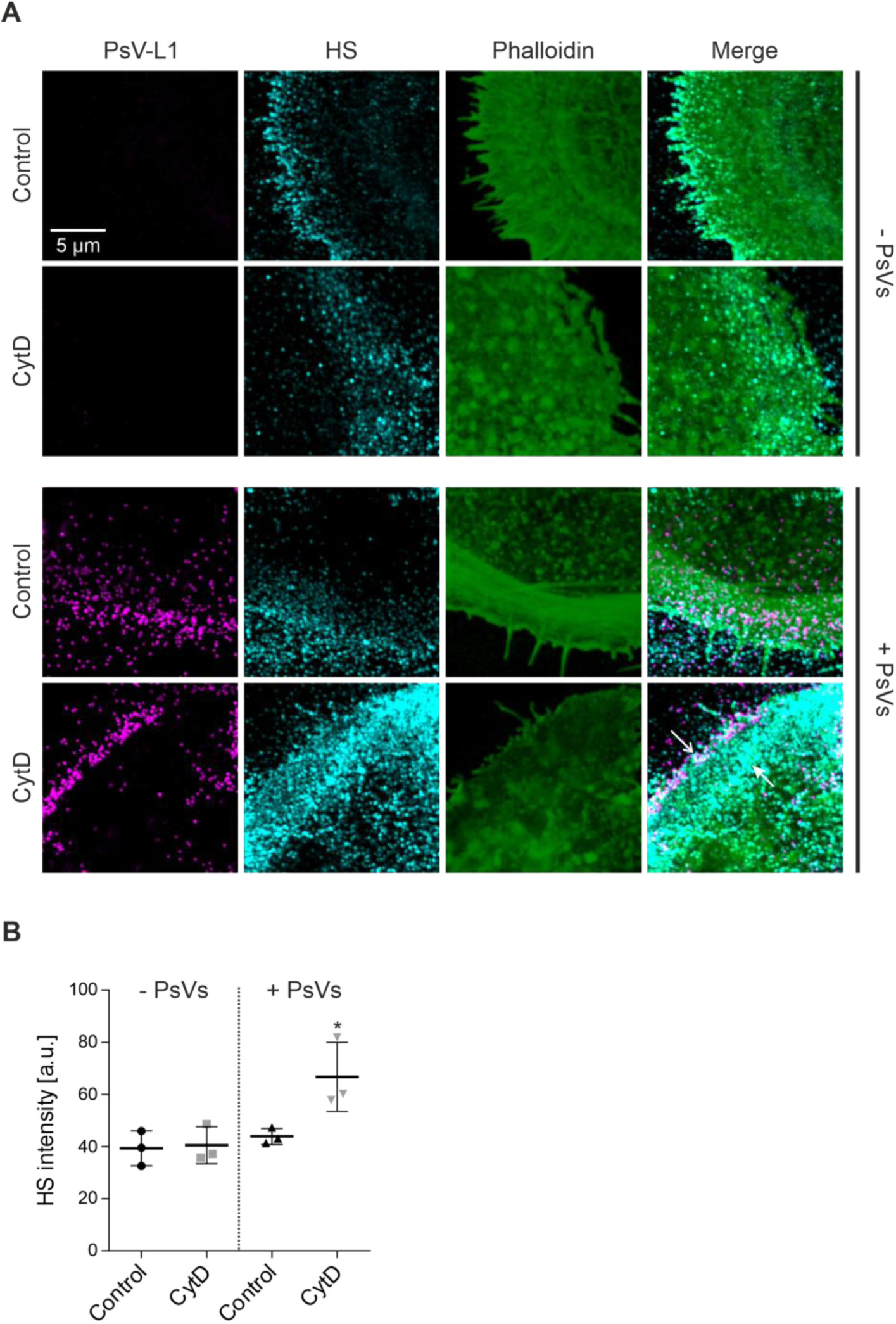
Increased HS intensity after incubation with PsVs and CytD. (A) HaCaT cells were incubated without (top) and with (bottom) PsVs at 37 °C for 5 h, in the absence (Control, upper panels) or presence of 10 µg/ml CytD (CytD, lower panels). Afterwards, cells were washed, fixed and stained. Immunofluorescence was used for L1 (STAR GREEN; magenta LUT) and for HS (AlexaFluor 594; cyan LUT) staining. F-actin was stained by iFluor647-labelled phalloidin (green LUT). PsVs-L1, HS and F-actin staining were imaged in the confocal mode of a STED microscope. The open arrow marks a region where PsVs overlap with HS. The closed arrow marks a region devoid of PsVs showing strong HS staining. (B) For analysis of the mean HS intensity, we placed large ROIs onto the images covering mainly the cell body but including parts of the cell periphery as well. Values are given as means ± SD (n = 3 biological replicates). Statistical differences between Control and CytD were analyzed by using the two-tailed, unpaired student’s *t*-test (n = 3, for details see methods).

Next, we used an antibody that reacts with a HS neo-epitope generated by heparitinase-treated heparan sulfate chains (***Yokoyama et al., 1999***; for details see methods). This neo-epitope staining is not affected by CytD and the incubation time (Supplementary Figure 1), suggesting that CytD does not directly affect HS processing. Collectively, our findings indicate that without actin-dependent PsV recruitment HS cleavage products are retained in the ECM, consistent with the hypothesis that cleaved HS remains associated with PsVs (***Ozbun and Campos, 2021***).

### Fast recruitment of PsVs to the cell body

Next, we studied the time course of PsV recruitment at higher time resolution. Cells were treated as in Figure 1A (in Figure 4, the 0 min time point is identical), followed by removal of PsVs/CytD. Different from Figure 2, we did not calculate the PCC that provides only an approximate idea of the time course, but monitored the diminishment of the integrated density of PsVs locating at the cell periphery, after 0, 15, 30 and 60 min. To delineate the cell periphery from the cell body, the TMA-DPH membrane staining is used as a reference (Figure 4A, the cell periphery is defined by the area enclosed by the two white lines in the PsV-DNA images (magenta LUT)).

**Figure 4.**
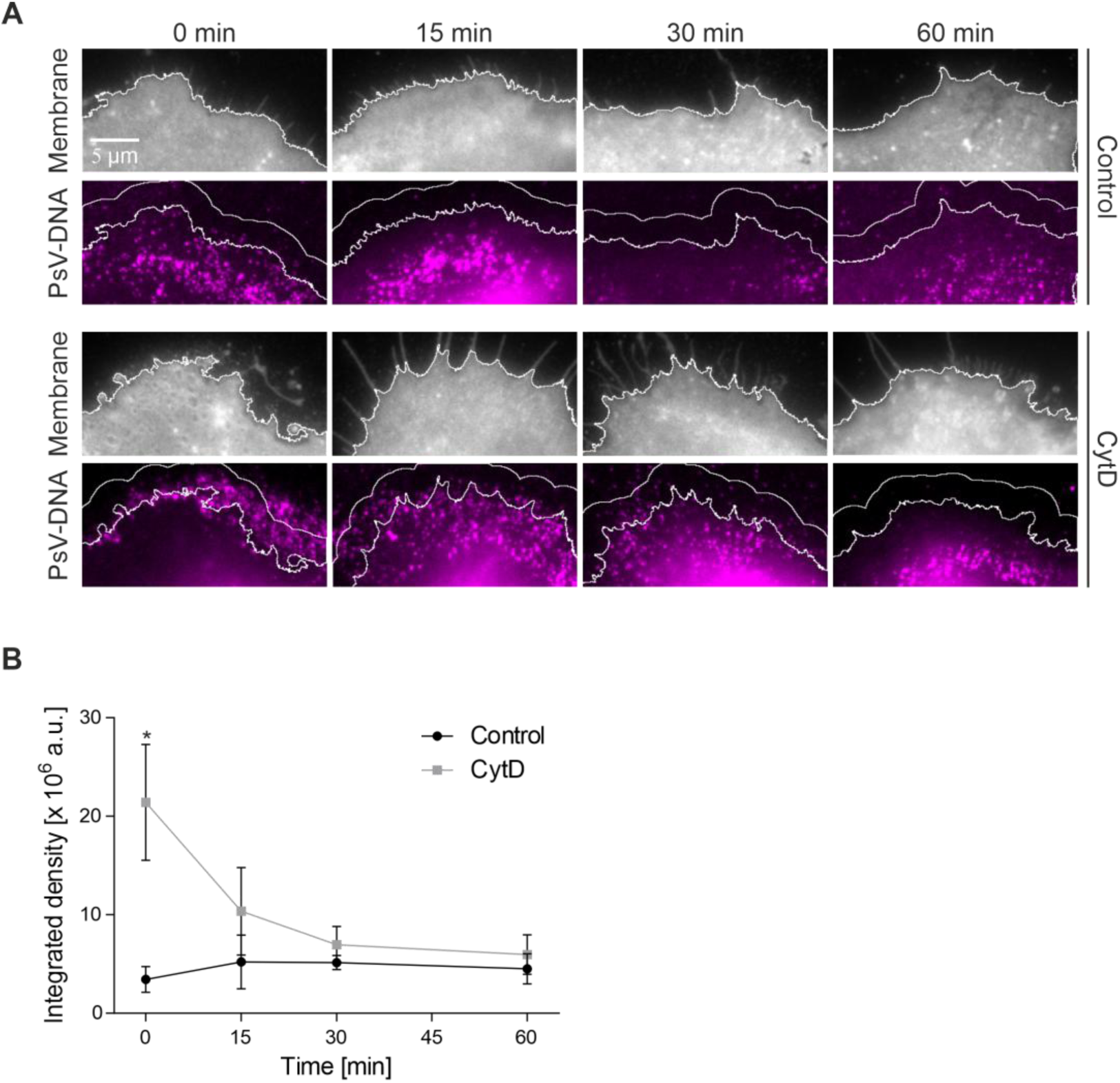
Fast diminishment of accumulated PsVs at the cell periphery after removal of CytD. (A) HaCaT cells were incubated with PsVs at 37 °C for 5 h, in the absence (Control) or presence of 10 µg/ml CytD (CytD). Then, cells were washed and incubated for the indicated time periods without PsVs/CytD, before they were fixed and stained as in Figure 1 (t = 0 min is identical to Figure 1A; for clarity we show only the membrane (gray LUT; images are shown at different settings of brightness and contrast) and the PsV-DNA staining (magenta LUT; images are shown at the same settings of brightness and contrast). The white lines in the membrane images delineate the cell body from the periphery. They were created with reference to the membrane staining (for details see methods). Using the cell body delineation as starting point, an up to 30-pixel broad area was created (PsV channel, magenta LUT; areas enclosed by the smoother white lines and the cell body delineation lines). The areas enclosed by the white lines define the cell peripheries. (B) The PsV-DNA signal of the periphery was quantified as integrated density, background corrected, and plotted over time. Values are given as means ± SD (n = 3 biological replicates). The statistical difference between the same time points of Control and CytD were analyzed by using the two-tailed, unpaired student’s *t*-test (n = 3, for details see methods).

At 0 min, compared to the control, CytD causes in the periphery a 6-fold increase of PsV signal (Figure 4B). This increase is more than halved when cells were incubated for 15 min, and after 30 min, the level of the control is reached (Figure 4B). This suggests that the half-time of PsV recruitment from the periphery to the cell body is about 15 min. Hence, active recruitment is fast and therefore cannot be a bottleneck in the time course of infection. We conclude that active recruitment is not responsible for the asynchronous virion uptake observed for HPV PsVs.

### Recruitment of PsVs to CD151

PsVs may approach the entry factor CD151 early, already during recruitment. For studying this possibility, we employ superresolution STED microscopy, analyzing the association of PsVs with CD151 over time. Cells are treated and monitored as above, but with an extended time window of up to 180 min, as cell surface processes are expected to take longer than an hour. After fixation, PsVs and CD151 were double-stained with antibodies against L1 (Figure 5A, magenta LUT) and CD151 (Figure 5A, green LUT). In addition, we stained F-actin with fluorescent labelled phalloidin (shown in Supplementary Figures 2 and 3). We simultaneously image L1 and F-actin in the confocal and CD151 in the STED channel.

**Figure 5.**
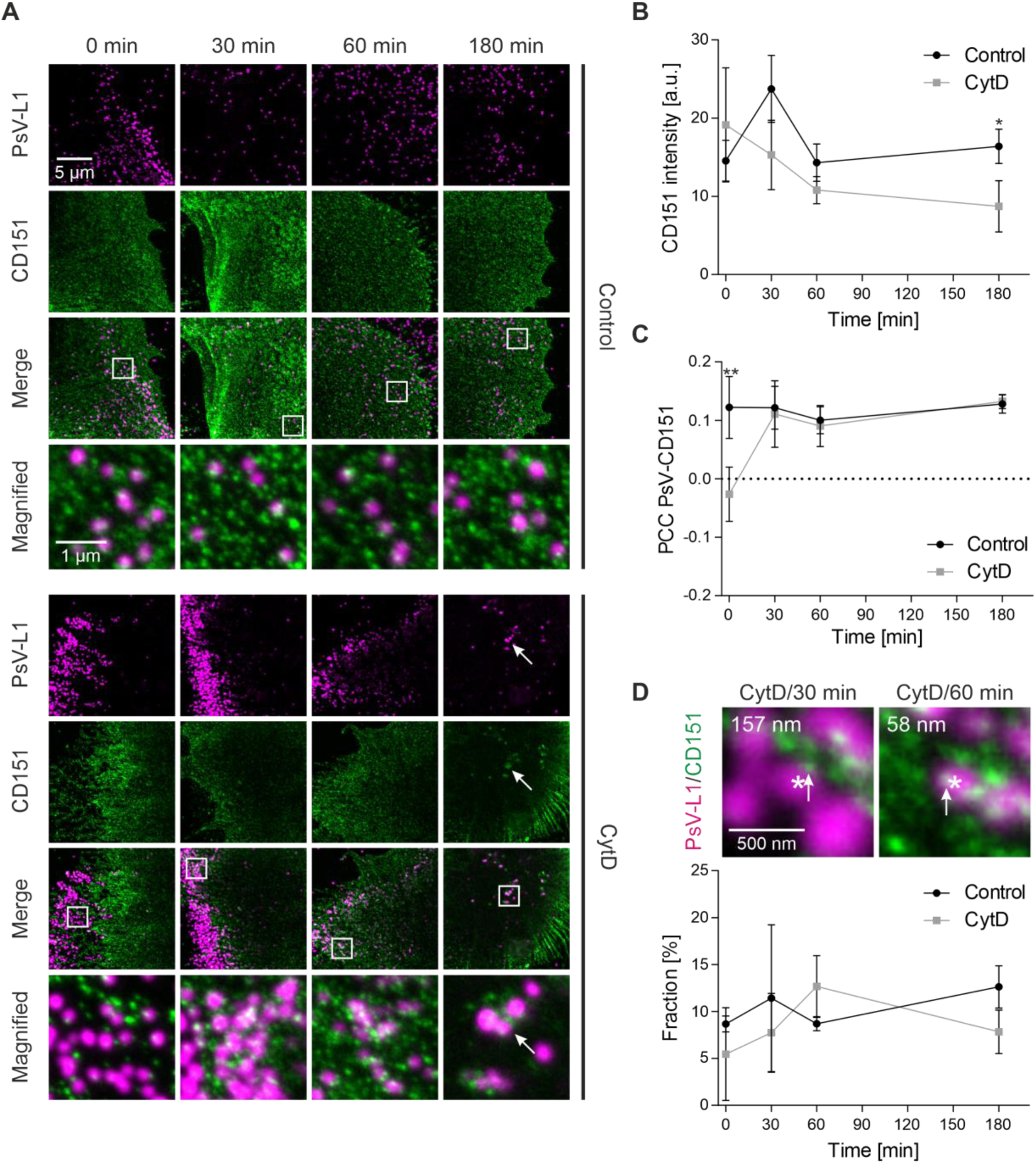
Association between PsVs and CD151 occurs early in the infection cascade. (A) HaCaT cells were incubated with PsVs at 37 °C for 5 h, in the absence (Control, upper panels) or presence of 10 µg/ml CytD (CytD, lower panels). Afterwards, cells were washed and incubated without PsVs/CytD further for 0 min, 30 min, 60 min or 180 min, before they were fixed and stained by indirect immunofluorescence for L1 (STAR GREEN; magenta LUT) and for CD151 (AlexaFluor 594; green LUT), and for F-actin by iFluor647-labelled phalloidin (here not shown for clarity, please see Supplementary Figure 2A for F-actin staining illustrating the variability of filopodia after CytD treatment). The bottom rows show magnified views of the merged images from the regions marked by the white boxes. PsV-L1 and F-actin staining were imaged in the confocal and CD151 staining in the STED mode of a STED microscope, respectively. Therefore, compared to CD151, the PsVs are less resolved and also appear much larger than their real physical size (see magnified views). CytD/180 min, arrows mark presumably endocytic structures that formed in the central cell body region (for more examples see Supplementary Figure 4). For analysis, we placed large ROIs onto the images that covered mainly the cell body but included parts of the cell periphery as well (for an example ROI see Supplementary Figure 5A). (B) The mean CD151 intensity was measured and plotted over time. (C) The PCC between PsV-L1 (magenta LUT) and CD151 (green LUT) was calculated and plotted over time. (D) The fraction of PsVs (in percent) that have a distance to the next neighbored CD151 maximum ≤ 80 nm, which we define as closely associated, is plotted over time. Please note that the values in (D) were corrected for random background association (for details see Supplementary Figure 7). Two examples of PsVs (each marked by an asterisk) from the CytD/30 min (left) and CytD/60 min (right) conditions are shown. The value in the upper left states the shortest distance between the PsV and the next nearest CD151 maximum (marked by an arrow) in nm. Values are given as means ± SD (n = 3 biological replicates). Statistical differences between the same time points of Control and CytD were analyzed by using the two-tailed, unpaired student’s *t*-test (n = 3, for details see methods).

As shown in Figure 5A, CD151 concentrates in spots scattered across the cell surface. It is also present at cell protrusions that are rich in F-actin, and that vary strongly in number and shape (Supplementary Figure 2A shows some examples of these filopodia). Compared to the control, CytD increases the occurrence of accumulated PsVs at the cell periphery, and at early time points PsVs appear brighter (Figure 5A; for overview images and F-actin staining of 0 min/CytD see Supplementary Figure 3A and B; for variability of PsV accumulations after CytD see Supplementary Figure 3B), as already observed in Figure 1B to D.

Initially, we wondered whether, after reaching the cell body edge, PsVs are transported quickly further towards the center. Based on the CD151 image, a cell border region was defined (for details see Supplementary Figure 2B). We counted the number of PsVs in this region, and expressed it as percentage of all PsVs in the image. In the control, the fraction of PsVs in the cell border region was rather low, ranging from 13.4% (0 min) to 7.6% (180 min). After CytD, at 0 min, compared to the control the fraction almost tripled (36.5%). It diminished to 16.9% after 60 min, and 12.3% after 180 min (Supplementary Figure 2C). This suggests that most PsVs leave the cell periphery within 1 h.

The intensity of the CD151 staining between cells is highly variable (Figure 5A and B). In the control, the mean CD151 intensity shows no trend over the 180 min time course, whereas with CytD the intensity after 180 min is diminished in comparison to the control (Figure 5B). The decrease of the CD151 staining intensity points towards the possibility that, after CytD wash off, CD151 is more strongly internalized compared to the control, presumably due to increased co-internalization with endocytosed PsVs. This idea is supported by the observation that, in particular at CytD/180 min, we occasionally observe CD151/PsV agglomerations (Figure 5A, see structures marked by arrows at CytD/180 min, for more examples see Supplementary Figure 4A). We did not study this issue systematically, but some of these structures have clear three-dimensional extension (see Supplementary Figure 4B for axial scans). Therefore, they are likely tubular structures filled with several PsVs, as previously described by electron microscopy (***Schelhaas et al., 2012***). We observe fewer of such structures at control/180 min, probably because cells have been actively interacting with PsVs for altogether 8 h, opposed to 3 h in the CytD treated cells. Hence, after CytD wash off, PsV endocytosis may be more synchronized compared to control, explaining the CD151 staining intensity diminishment at CytD/180 min (Figure 5B).

For studying the association between PsVs (L1) and CD151, the PCC between the channels is calculated. The PCCs are around 0.1, with the exception of the 0 min/CytD value that is significantly lower and even negative (Figure 5C, for PCC values of flipped images see Supplementary Figure 5B and C). This reflects the in part mutual exclusion of the two stainings as already discussed above; PsV accumulations tend to be at the cell periphery and the CD151 staining is at the cell body and in filopodia. In any case, the lower PCC at 0 min/CytD suggests that without active recruitment less PsVs reach CD151. At 30 min after CytD, the PCC has reached the level of 0.1 as in the control, which is in line with the idea of fast recruitment as observed in Figure 4. To follow how the basal cell membrane is populated with PsVs over time, as additional analysis we determined the PsVs per µm^2^ in ROIs placed in the cell body region. At 0 min, CytD reduces the PsV density to 19 - 33%, albeit the effect is not significant, and at 180 min/CytD the same PsV density as in the control is reached (Supplementary Figure 6A and B). Overall, under CytD there was a trend towards less PsVs present (Supplementary Figure 6A and B). Hence, both Figure 5C and Supplementary Figure 6A and B suggest that active virion transport is required to reach efficiently the basal membrane.

We next studied the fraction of PsVs that are closely associated with CD151. As criteria for close association, we define a distance of ≤ 80 nm between PsV and CD151 maxima because this value is close to the resolution limit of the used microscope (***Finke et al., 2020***). In the control, the fraction of PsVs closely associated with CD151 is around 10% (Figure 5D, control), after correction for random background association, for which we used a calibration line based on the same density of PsVs in flipped images (see Supplementary Figure 7). At 0 min after CytD, we start with a fraction of 5.5% of closely associated PsVs, that increases to 12.7% in the next 60 min, although this increase is not significant (Figure 5D, CytD). In any case, it is remarkable that in the absence of active recruitment (0 min/CytD) we find half the fraction of closely associates PsVs compared to the control. This indicates that PsVs and CD151 associate very early in the infection cascade, essentially in the moment the PsVs reach the edge of the cell body.

In summary, we conclude that within 180 min after a 5 h pre-incubation with PsVs (Figure 5D, control), we encounter a type of steady-state situation (new PsVs are recruited whereas older ones disappear by endocytosis), in which about 10% of the PsVs are associated closely with CD151. CytD diminishes only the PsV-CD151 association at 0 min (5.5%, Figure 5D, CytD), suggesting that PsVs establish contact to CD151 early (see also the increase in the PCC between 0 min/CytD and 30 min/CytD, Figure 5C).

### PsV association with HS

Next, we studied the association between PsVs and HS. As a reference staining for the cell body, we used Itgα6 that is not visible at cell protrusions. PsVs are visualized by click-chemistry and imaged in the confocal channel. HS and Itgα6 were stained by antibodies, and imaged at STED microscopic resolution. The three stainings were simultaneously recorded.

As shown in Figure 6A (green LUT), the Itgα6 staining results in a pattern of dense spots. The Itgα6 intensity does not change over time (Supplementary Figure 8E). The pattern of the HS staining (cyan LUT) and the overlap of HS with PsVs and Itgα6 are highly variable (Figure 6A).

**Figure 6.**
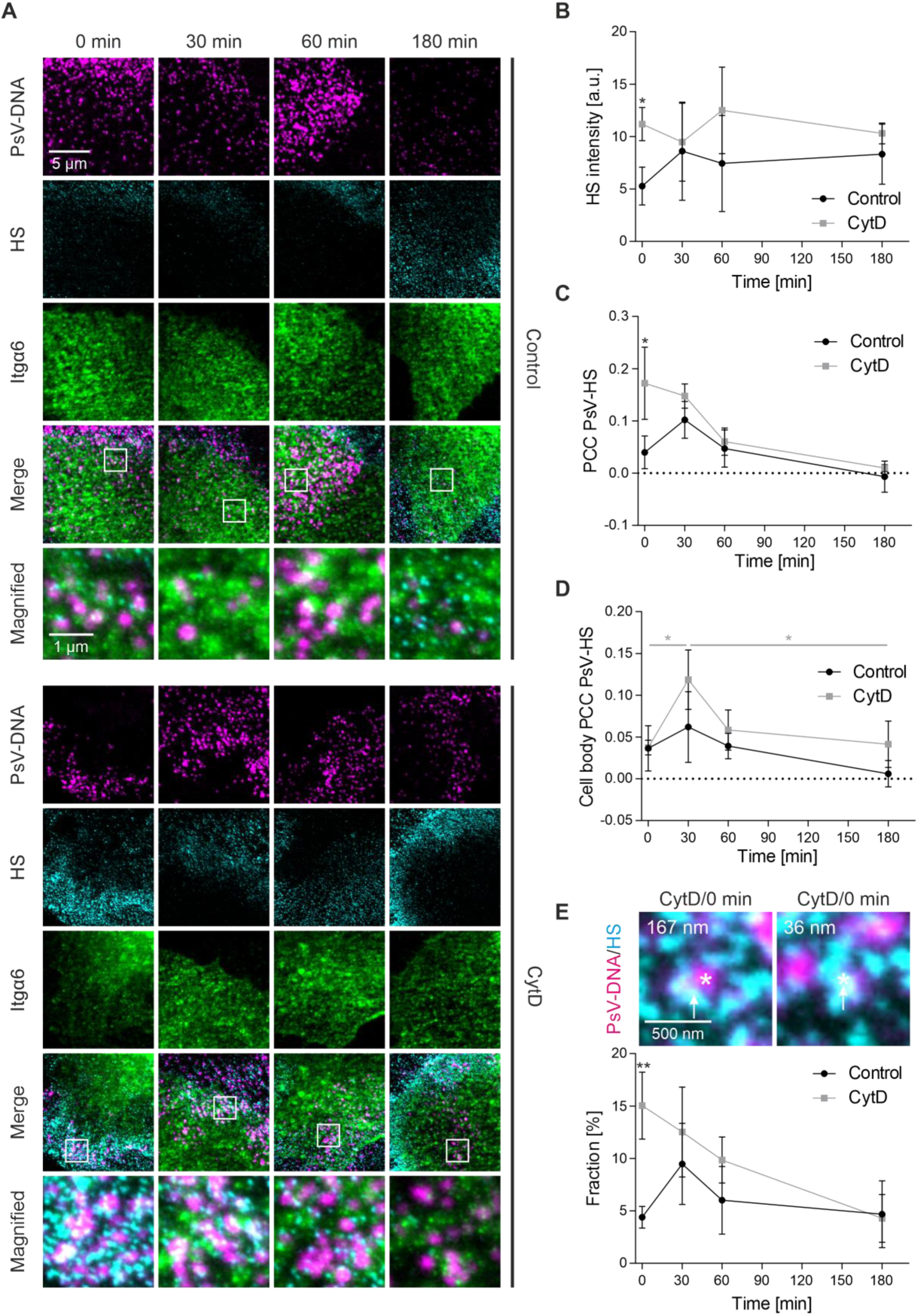
Association between PsVs and HS after CytD treatment. (A) HaCaT cells were incubated with PsVs at 37 °C for 5 h, in the absence (Control, upper panels) or presence of 10 µg/ml CytD (CytD, lower panels). Afterwards, cells were washed and incubated without PsVs/CytD further for up to 180 min, before they were fixed and stained. PsVs (magenta LUT) were visualized by click-chemistry (6-FAM Azide) and indirect immunolabeling was used for HS (AlexaFluor 594; cyan LUT) and for Itgα6 (STAR RED; green LUT). The bottom rows show magnified views of the white boxes in the merged images. PsV-DNA staining was imaged in the confocal and HS and Itgα6 staining in the STED mode of a STED microscope. For analysis, we placed large ROIs onto the images that covered mainly the cell body but included parts of the cell periphery as well (see example in Supplementary Figure 8A). For (D), smaller ROIs covering only the cell body region were used. (B) The mean HS intensity plotted over time. (C) The PCC between PsV-DNA (magenta LUT) and HS (cyan LUT) over time. (D) The PCC between PsV-DNA (magenta LUT) and HS (cyan LUT) in the region of the cell body over time. (E) The fraction of PsVs (in percent) closely associating with HS (distance ≤ 80 nm) plotted over time (for background correction see Supplementary Figure 9). Two examples of PsVs (each marked by an asterisk) from the CytD/0 min condition are shown. The value in the upper left corner states the shortest distance (in nm) between the marked PsV and its next nearest HS maximum (marked by an arrow). Values are given as means ± SD (n = 3 biological replicates). Using the two-tailed, unpaired student’s *t*-test (n = 3 biological replicates), we analyzed in (B), (C) and (E) the statistical differences between the same time points of Control and CytD, and in (D) the difference between CytD/30 min and CytD/0 min or CytD/180 min (for details see methods).

CytD increases the intensity of HS (Figure 6B; also apparent when comparing the HS brightness in the upper and lower panels of Figure 6A; for an overview of the CytD/0min images see Supplementary Figure 3C). This increase of intensity is particularly notable at the 0 min time point, where the samples treated with CytD have a more than two-fold higher intensity and differ significantly from the control. Hence, in this experiment we reproduce the PsV/CytD mediated HS intensity increase observed in Figure 3B.

In the control, the PCC between PsVs and HS are low with no clear trend. The largest PCC is found at CytD/0 min, which reflects the finding that at this time point both PsVs and HS preferentially locate at the cell periphery (Figure 6C, for PCC values of flipped images see Supplementary Figure 8B and C). Over time, the accumulated PsVs diminish due to recruitment, which is accompanied by a PCC approaching zero, as in the control (Figure 6C, 180 min). Additionally, we analyze the PCC between PsVs and HS specifically in the cell body region, excluding the cell periphery (Figure 6D). For the control, the PCC in the cell body region ranges between 0 and 0.05. After CytD, we observe an increase in the PCC from 0 min to 30 min and a decrease from 30 min to 180 min (for PCC values of flipped images see Supplementary Figure 8F and G). When analyzing the PsVs closely associating with HS, at 0 min, CytD increases the fraction of PsVs associated with HS more than 3-fold. Over the next 180 min, this fraction gradually decreases until it is equal to the control value (Figure 6E, 180 min).

Altogether, the analysis of the PCC between HS and PsVs (Figure 6C) along with the fraction of closely HS associated PsVs (Figure 6E) indicates that CytD treatment increases HS/PsV association at 0 min, likely because PsVs remain accumulated in the ECM. After CytD removal, this association diminishes. An increase in PCC is observed at 30 min specifically in the region of the cell body (Figure 6D) which likely reflects the recruitment of HS-coated virions to the cell body. However, as this association diminishes over the subsequent 150 min, an increasing fraction of PsVs is no longer associated with HS due to progressive loss of the HS coat over time, or alternatively, PsVs just may disappear after internalization.

Next, for each PsV we plot the ‘distance to the next nearest Itgα6 maximum’ against ‘the distance to the next nearest HS maximum’ (Figure 7). In the control, the distances remain essentially unchanged over the entire observation time; 59% - 65% of the PsVs are at a short distance (< 250 nm) to both Itgα6 and HS (Figure 7B, black; see also Supplementary Table 1), 12 - 15% are at a short distance (< 250 nm) to HS but not to Itgα6 (> 250 nm) (Figure 7B, green; see also dashed green box in Figure 9 marking PsVs in the ECM). Regarding the distance plots of untreated cells in Figure 7 (please compare the upper distance plots from left to right), they suggest that the distance patterns do not change over the 180 min observation time. In cells treated with CytD, a larger fraction of PsVs is accumulated at the periphery at the 0 min time point, which is reflected in a larger fraction (46% as opposed to 12% in the control; Figure 7B, green; Supplementary Table 1) of PsVs with a short distance (< 250 nm) to HS but not to Itgα6. In the distance plots, it can be noticed that the PsVs in the CytD treated cells over time acquire shorter distances to Itgα6 and larger distances to HS (Figure 7C, lower row). The population with short distances to HS and large ones to Itgα6 strongly diminishes from 46% to 11% (Figure 7B, CytD, green). After 180 min, the distances are similar to the untreated control. Between CytD/60 min and CytD/180 min, the fraction of PsVs with a large distance (> 250 nm) to HS and a short distance (< 250 nm) to Itgα6, representing PsVs at the cell body without HS (see also dashed magenta box in Figure 9), increases from 9.6% to 25.2% (Figure 7B, magenta). This is inconsistent with the idea of endocytosis of HS-coated PsVs that would only diminish short distances to Itgα6 but not create long distances to HS, which is what we observe. The observation supports the idea that PsVs lose their HS coat after translocating to the cell surface, which is in line with the transient increase in the cell body PCC between HS and PsVs at 30 min (Figure 6D).

**Figure 7.**
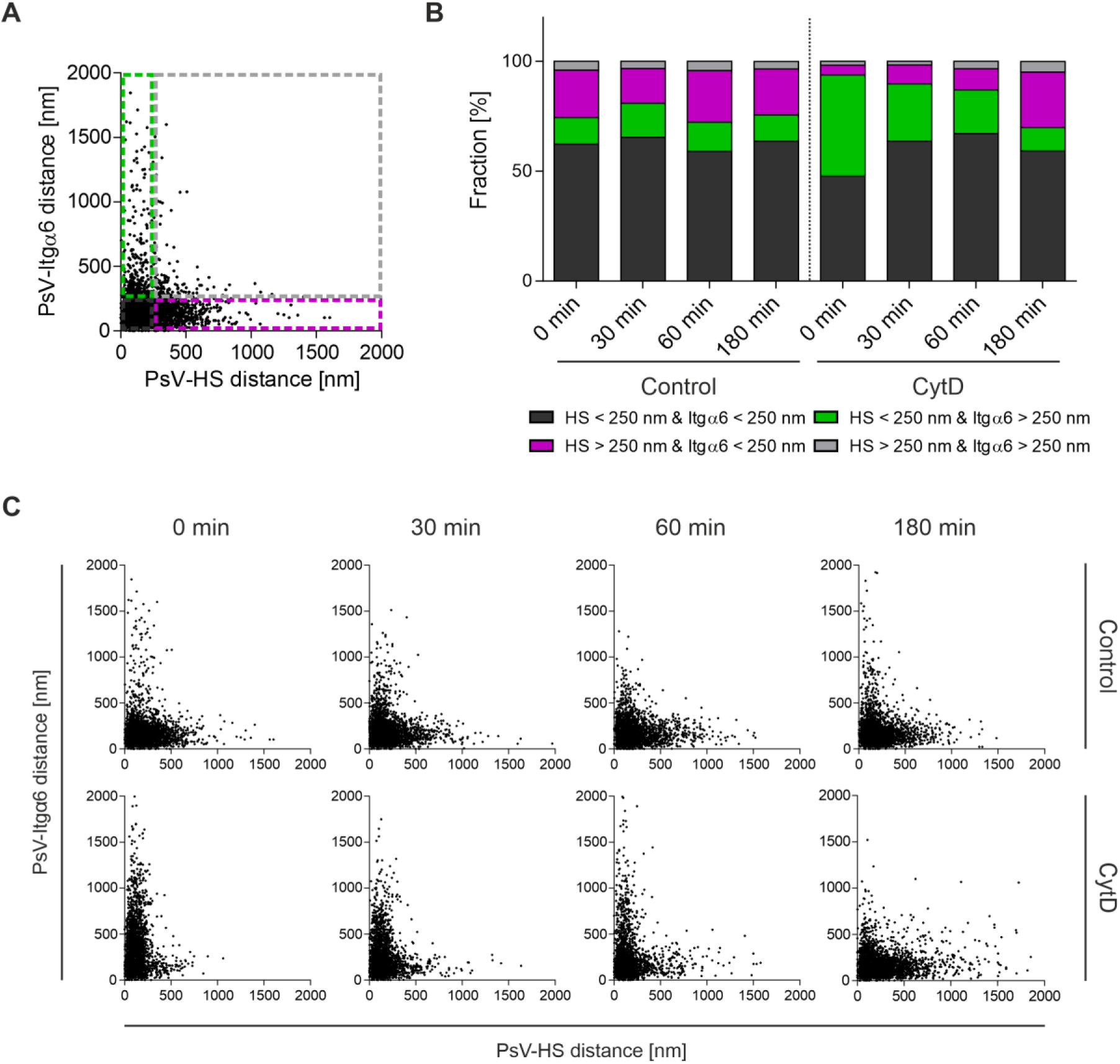
PsV-Itgα6 and PsV-HS distances over time. (A) Definition of four PsV populations based on the PsV distances to Itgα6 and HS (please note that the data does not include distances of exactly 250 nm wherefore symbols as ≥ and ≤ are omitted; the plot is taken from Control, 0 min and shown again in C). Dashed green rectangle, PsVs with a distance to HS < 250 nm and to Itgα6 > 250 nm. Dashed magenta rectangle, PsVs with a distance to HS > 250 nm and to Itgα6 < 250 nm. Dashed gray square, PsVs with a distance to HS > 250 nm and to Itgα6 > 250 nm. PsVs not included in the previous categories have a distance to HS < 250 nm and to Itgα6 < 250 nm (dashed black square). (B) From the PsVs analyzed in Figure 6, for the Control (left) and CytD (right) the PsV fraction size (in percent) of each population is illustrated. Shown are the means of three biological replicates. For means ± SD and statistical analysis see Supplementary Table 1. (C) For the Control (top) and CytD (bottom), for each PsV, we plotted the shortest distance to Itgα6 against the shortest distance to HS (pooling the three biological replicates; 3,043 – 4,080 PsVs per plot).

The recruitment of HS towards the cell body after removal of CytD, which indirectly demonstrates that PsVs are coated with HS, is suggested by a shortening of the HS-Itgα6 distance over 180 min only after CytD wash off (Supplementary Figure 8D). Together with the observation that the PCC between PsVs and HS in the cell body area increases (Figure 6D), the data suggest that HS-coated PsVs are recruited to the cell body. A fraction of PsVs sheds HS within one hour after removing CytD (Figure 7C, lower panels, showing increasing PsV-HS distances after 60 min). This is the time window in which we notice the occurrence of endocytic PsV/CD151 structures (see above and Supplementary Figure 4B).

Actin retrograde transport underlies the filopodial virion transport and is the integrative result of three components (***Smith et al., 2008; Schelhaas et al., 2008***). On the one side, actin filaments in the cell periphery are pushed into the cell body area when actin polymerizes at their tips. The retrograde filament movement causes tension and is facilitated by F-actin degradation opposed to the side of F-actin growth. Moreover, the motor protein myosin II exerts force on actin filaments, pulling those towards the cell body. As CytD broadly interferes with F-actin dependent processes like filopodial transport and cell migration, we investigated the effects upon inhibition of only one component of actin retrograde flow, namely the myosin II mediated retrograde movement towards the cell body. Instead of CytD, we employed in the 5 h preincubation the myosin II inhibitor blebbistatin (***Schelhaas et al., 2008***). For the control (0 min), we show in Figure 8A one example of a cell with comparatively many PsVs at the periphery (as mentioned above, the PsV pattern is highly variable and also in the control we occasionally observe PsV accumulations) to better illustrate the difference to the PsV pattern occasionally seen with blebbistatin. After blebbistatin treatment (0 min), PsVs do not reach the central cell body as in the control, but are less dispersed than after CytD treatment, seemingly as if recruitment started but stopped in the midst of the pathway (Figure 8A, blebbistatin). The PCC between PsVs and HS, like after CytD (Figure 6C), is elevated after blebbistatin, albeit the effect is not significant (Figure 8C). The cell body PCC is not, as under CytD, at 30 min (Figure 6D) but already at 0 min elevated (compare Figure 6D to Figure 8D), which can be explained by partial recruitment. This is further supported by the fact that only 8% of PsVs are closely associated with HS (Figure 8E; blebbistatin, 0 min) compared to 15% after CytD treatment (Figure 6E; 0 min). Furthermore, after 0 min PsV incubation with blebbistatin we observe no effect on the HS intensity (compare Figure 8B to Figure 3B and Figure 6B). Hence, in contrast to CytD, blebbistatin does not preserve the PsVs in the ECM where they associate with HS, but ongoing actin polymerization may push actin filaments along with PsVs towards the cell body.

**Figure 8.**
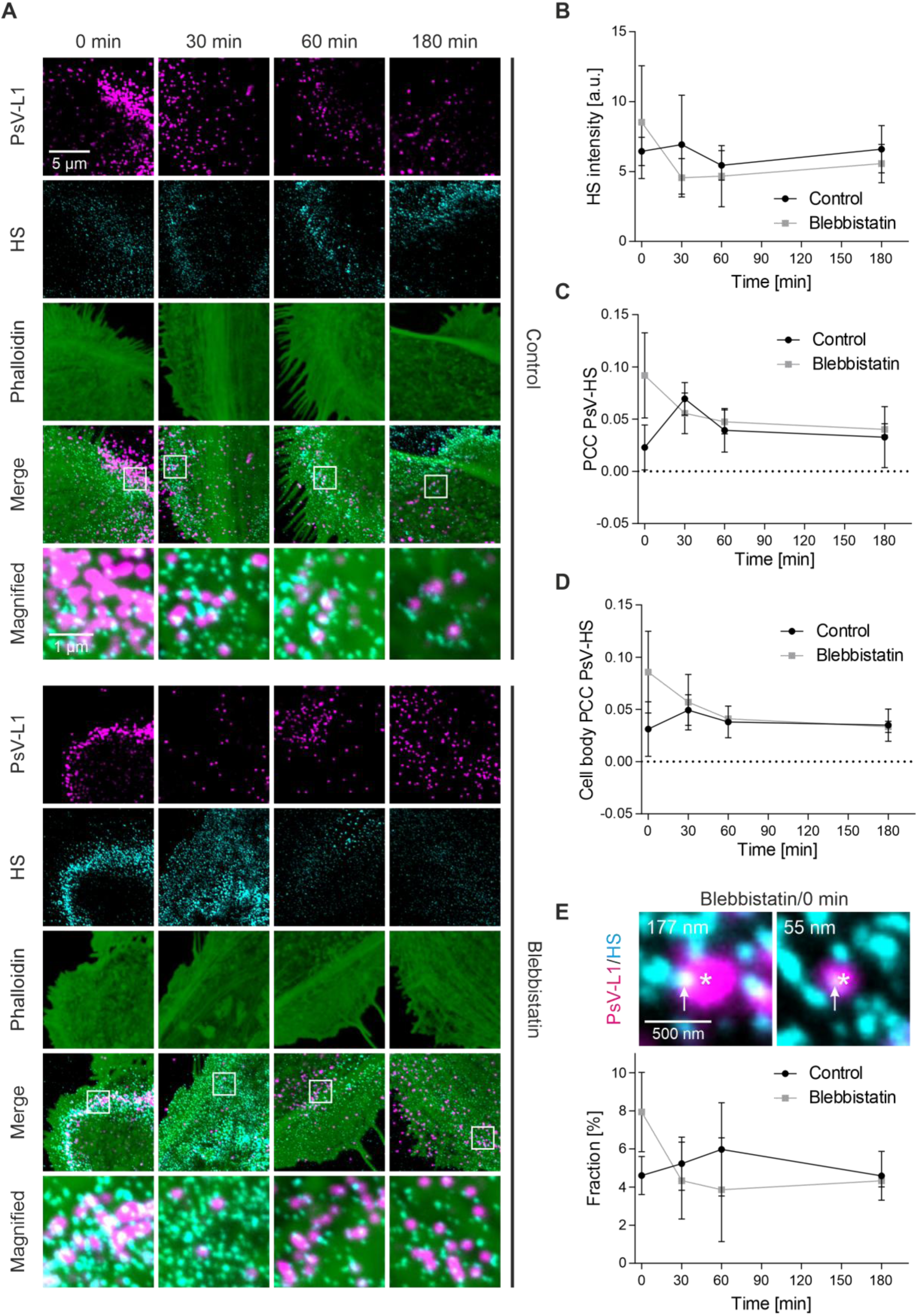
Association between PsVs and HS after blebbistatin treatment. (A) HaCaT cells were incubated with PsVs at 37 °C for 5 h, in the absence (Control, upper panels) or presence of 30 µM blebbistatin (Blebbistatin, lower panels). Afterwards, cells were washed and incubated without PsVs/blebbistatin further for up to 180 min, before they were fixed and stained. Immunofluorescence was used for L1 (STAR GREEN; magenta LUT) and HS (AlexaFluor 594; cyan LUT) staining. F-actin was stained by iFluor647-labelled phalloidin (green LUT). The bottom rows show magnified views of the white boxes in the merged images. PsVs and F-actin staining were imaged in the confocal and HS staining in the STED mode of a STED microscope. For analysis, we placed large ROIs onto the images that covered mainly the cell body but included parts of the cell periphery as well. For (D), smaller ROIs covering only the cell body region were used. (B) Mean HS intensity over time. (C) PCC between PsV-L1 (magenta LUT) and HS (cyan LUT) over time (for control with flipped images see Supplementary Figure 10A and B). (D) PCC between PsV-L1 (magenta LUT) and HS (cyan LUT) in the region of the cell body over time (for control with flipped images see Supplementary Figure 10C and D). (E) The fraction of PsVs (in percent) closely associating with HS (distance ≤ 80 nm) plotted over time (for background correction see Supplementary Figure 11). Two examples of PsVs (each marked by an asterisk) from the blebbistatin/0 min condition are shown. The value in the upper left states the shortest distance between the PsV and its next nearest HS maximum (marked by an arrow) in nm. Values are given as means ± SD (n = 3 biological replicates). Statistical differences between the same time points of Control and Blebbistatin were analyzed by using the two-tailed, unpaired student’s *t*-test (n = 3), but the analysis yielded no p-values below 0.05.

### PsV binding in the absence of a diffusion barrier

Throughout all experiments, we observe at 0 min/CytD only few PsVs at the basal membrane (Figure 1A, Supplementary Figure 6A and B; see also PCC at 0 min between PsVs an CD151 in Figure 5C), suggesting that in the absence of active recruitment the access to the basal membrane via passive diffusion is limited. We wondered, how many PsVs may bind to the cell membrane without a diffusion barrier? For this reason, we incubated EDTA detached HaCaT cells in suspension with PsVs for 1 h at 4 °C, followed by re-attachment for 1 h. Under these conditions, we find, despite of a shorter incubation time (1 h versus 5 h), a roughly 3-fold larger PsV density (1.7 PsVs/µm^2^ (Supplementary Figure 6D)) than the highest density observed in the other experiments. However, it should be noted that values of the different experiments cannot be directly compared. Aside from the different treatments, another difference lies in the size of the imaged membrane. The re-attachment of cells is not complete after 1 h (compare size of adhered membranes in Supplementary Figure 6A and 1A), wherefore the membranes are likely strongly ruffled, which results in the underestimation of the membrane area. As a result, we overestimate the PsVs per µm^2^ adhered membrane (please note that we cannot re-attach cells for longer times as we then lose PsVs due to endocytosis). In any case, the experiment suggests that PsVs bind more efficiently to membrane surface receptors without a diffusion barrier. We conclude that in our assay PsVs cannot readily bypass the active PsV recruitment by diffusing directly to the basal cell membrane, which is plausible, because to make this happen a 55 nm large PsV must diffuse through the narrow gap between glass-coverslip and adhered cell.

## Discussion

In this study, we investigate the early events in HPV16 infection occurring at the cell surface. Our findings reveal that in our assay actin dependent mechanisms, rather than passive virion diffusion, are crucial for recruiting HS-coated virus particles from the adhesive ECM to the basal cell membrane. PsVs associate early with CD151, and shed their HS-coat before endocytosis. Moreover, we propose that reversible blocking of active recruitment by CytD allows for a more synchronized observation of the steps on the cell surface.

### A reversible CytD blocking of PsV recruitment from the ECM to the basal cell membrane

Many viruses use cell surface HS for primary attachment, including herpes simplex virus type 1, human cytomegalovirus, human immunodeficiency virus type 1, adenovirus type 2, dengue virus, hepatitis B virus, and vaccinia virus (***Cagno et al., 2019; Giroglou et al., 2001; Tian et al., 2021***).

HPVs could be different as they bind as well to HS of the extracellular basement membrane, at least *in vivo* during the wounding and healing processes required for infection (***Day and Schelhaas, 2014; Kines et al., 2009; Ozbun and Campos, 2021; Schiller et al., 2010***).

By treating cells with CytD, we block actin-mediated recruitment, leading to a 6-fold accumulation of PsVs at the cell periphery after 5 hours (Figure 4B). After CytD removal, infectivity is reduced by 29% (Figure 1E). Some reduction is to be expected, as PsVs have ≈20% less time to complete infection as compared to cells that were not treated with CytD. Hence, the infection assay suggests that the treatment is largely reversible and only slightly harmful, if at all. However, the luciferase infection assay does not distinguish between actively recruited PsVs and PsVs that bind passively by diffusion to the upper membrane. The latter fraction likely dominates the total infection rate and should be less affected by CytD than the fraction of actively recruited PsVs. Therefore, if the infection pathway of a small fraction of actively recruited PsVs is irreversibly inhibited, we may not be able to detect this effect on the background of unaffected passively binding PsV.

Upon CytD washout, PsVs approach the basal membrane with a half-time of ≈15 minutes, as suggested by the diminishment of accumulated PsVs at the periphery (Figure 4B). The half-time could be longer if cell spreading is also underlying the translocation of PsVs onto the cell body. However, we assume that this is rather unlikely, as cell spreading would increase the PCC between PsVs and F-actin under a condition where PsVs are not-primed (and therefore not actively recruited) but cell spreading occurs, which is not the case in Figure 2B and D (CytD/leupeptin).

In Figure 4, we observe fast diminishment of the peripheral PsVs within 15 min. It is in principle possible that after CytD removal ECM-accumulated PsVs are merely washed off. However, after only 15 min, the effect of this could not have been very strong, considering the still 6-fold accumulation of PsVs after the 5 h incubation with CytD.

Furthermore, CytD pre-treatment appears to synchronize uptake: We observe endocytic structures after CytD treatment more frequently and the CD151 intensity diminishes over time only after the CytD treatment (Figure 5B). We estimate that, compared to control, about 6-fold more PsVs (Figure 4B) approach the cell surface in a coordinated fashion after CytD removal, which likely enhances the number of endocytic events compared to untreated cells. We therefore propose that the processing of the PsVs in the ECM is contributing to desynchronization and is largely completed after the 5 h pre-incubation with CytD.

### The role of actin-driven virion transport

The strong electrostatic interaction between HPV and HS implies that some processing — via HS cleavage or structural capsid changes — is a prerequisite for the release of virions from the ECM.

CytD likely does not inhibit HS processing (***Jamieson et al., 2021***) nor is it likely to inhibit KLK8. Therefore, we assume that, although PsVs are primed after 5 h of incubation with CytD, they remain in the ECM. They may be associated via weaker interactions, and actin-driven transport is required for pulling the HS-coated virions out of the sticky matrix towards the cell body. Sticky PsV aggregations that are released from HeLa cell surface after binding and blocking of the subsequent entry pathway have been observed earlier and might reflect similar HS/PsV-stuctures (***Mikuličić et al., 2025***). It is possible that an ECM bound virion is not exposed to all required processing factors at the same time. In this case, migration through the ECM enables for contact with all processing enzymes. Once the fully processed virions have reached the cell surface, detachment from the ECM may be facilitated by new bonds established between the PsVs and cell surface receptors. Now, the PsVs and cell surface receptors move towards the center of the cell body by intracellular dynamics which results in CD151 accumulation at virus binding sites (see Figure 5 as well as Supplementary Figure 4) and ultimately endocytosis.

Both CytD and blebbistatin increase PsV-HS colocalization, but blebbistatin allows partial recruitment (compare Figure 6C and Figure 8C as well as Figure 6D and Figure 8D), indicating that actin retrograde flow initiates movement, while myosin II contributes to sustained transport. As a result, virus movement can proceed to some extent after blebbistatin treatment. However, the subsequent blocking of transport at later time points reflects the inhibition of myosin II, which eventually disrupts the cytoskeletal dynamics and force generation that is necessary for sustained transport. In contrast, CytD directly interferes with actin polymerization, leading to an earlier and more persistent blocking of virus transport due to structural disruption of the filopodial tracks themselves.

Altogether, our findings support earlier reports showing that HPV utilizes filopodia for cell entry, migrating at several micrometers per minute (***Schelhaas et al., 2008; Smith et al., 2008***). This translocation is fast compared to the entire infection process and therefore cannot largely contribute to the asynchronous HPV uptake.

The analyzed PsVs hardly bind to the basal cell surface directly by diffusion (Supplementary Figure 6, compare PsV maxima density at 0 min/CytD in A and B to C). Therefore, the actin-driven virion transport would play a decisive role in HPV infection if cells would form a monolayer with a disruption at which ECM is present and that is approached by PsVs, a scenario similar to *in vivo* infection. In addition, cell migration could establish contact between PsVs and the cell surface.

### HaCaT cells as model system in HPV infection

Inhibition of HPV16 PsV transport diminishes infection only of subconfluent but not of confluent HaCaT cells by about 50% (***Schelhaas et al., 2008***). Therefore, actin-dependent translocation along protrusions may be dispensable for infection at a high cell density (***Schelhaas et al., 2008***) and merely increases the exposure of cells to virions (***Smith et al., 2008***) that can readily bind to the upper cell membrane. We are not aware of a PsV translocation mechanism from the upper to the basal membrane. Therefore, in our assay, PsVs bound to the upper membrane are not expected to show up at the basal membrane. Comparing 0 min of control and CytD (Supplementary Figure 6A and B), we find that compared to the control 19 - 33% of the PsVs reach the basal membrane in the absence of active transport, or in other words, most likely by passive diffusion. Actually, the range from 19 – 33% must be a strong overestimate as PsVs in the control are in transit and many actively recruited PsVs are already internalized during the 5 h incubation period. For this reason, we propose that most likely much less than 10% of the PsVs reach the basal membrane by diffusion. Moreover, in the absence of the diffusion barrier, the density of bound PsVs is strongly increased (Supplementary Figure 6D), showing indirectly that at the basal membrane the binding sites are difficult to access without active recruitment. Taken together, we propose the large majority of PsVs analyzed in our assay are ECM bound and actively recruited to the basal cell membrane.

The increased virion exposure via active recruitment could be relevant *in vivo*, as wounding of the epidermis results in upregulation of filopodia formation (***Vasioukhin et al., 2000***). Filopodia usage not only facilitates infection but also increases the likelihood of virions to reach their target cells during wound healing, namely the filopodia-rich basal dividing cells. In fact, several types of viruses exploit filopodia during virus entry (***Chang et al., 2016***), hinting at the possibility that for HPV and other types of viruses actin-driven virion transport may play a more important role than it is currently assumed. In this scenario, sub-confluent HaCaT cells, or even better single HaCaT cells, would be an ideal model system for the microscopic study of these very early infection steps that involve ECM attachment and subsequent active recruitment, as supposed to occur during *in vivo* infection of basal keratinocytes after binding of virions to the basement membrane (***Bienkowska-Haba et al., 2018; Day and Schelhaas, 2014; Kines et al., 2009; Schiller et al., 2010***). In contrast, in biochemical infection assays, virions diffusing to HSPGs on the cell surface, and by this bypassing active recruitment, are assayed together with the actively recruited virions. Should cells secrete little ECM and are grown to confluency, the passively binding virions are supposed to strongly dominate the infection rate in a biochemical infection assay.

### Recruited PsVs are decorated with HS which they lose over time

CytD treatment increases the HS signal when PsVs are present (Figure 3B and 6B). A relationship between PsVs and HS cleavage products has been previously suggested (***Ozbun and Campos, 2021; Surviladze et al., 2012***), and we propose that our data is in line with these previous observations. It should be noted that under conditions without the 5 h CytD blocking, PsVs may only briefly reside in the ECM and as a result are less decorated with HS. Hence, under our assay conditions the HS decoration of virions may be enhanced in comparison to other assays.

At the 0 min time point post-CytD, 15% of the PsVs are closely associated with HS, but this fraction declines to 4.3% within 180 min (Figure 6E). Considering the important role of HS as a primary attachment site, the fraction of 15% appears unexpectedly small. There are multiple possible explanations for underestimating the number of PsVs closely associated with HS. One factor may be that accumulated PsVs are not well-resolved (see above), which results in an underestimation of HS associated PsVs in particular at CytD/0 min. Other explanations include that the HS antibody does not reach all HS epitopes in the dense matrix. Yet another explanation could be that PsV binding to HS and antibody binding to HS is sometimes mutually exclusive, a hypothesis which is supported by data showing that the HPV capsid possesses multiple HS binding sites (***Dasgupta et al., 2011; Richards et al., 2013***). Finally, not all PsVs bound to the ECM are expected to bind to HS but could also bind to laminin 332 (***Culp et al., 2006***). Together, our data suggests that PsVs initially are recruited with HS attached to them and then shed it, potentially either upon receptor binding or further capsid rearrangement. These observations are in line with a previous study showing reduced HS colocalization during productive infection (***Selinka et al., 2007***).

The initial HS-coat may facilitate the formation of a complex with PsVs and cell surface receptors such as integrins or growth factor receptors (GFRs) (***Ballut et al., 2013; Pellegrini, 2001***). This could facilitate receptor clustering and engagement as shown previously for coronavirus infection (***Bugatti et al., 2025; Clausen et al., 2020; Zhang et al., 2023***), as well as the binding of signaling molecules to the cell surface (***Miladinova et al., 2022***). Here, CD151 may support receptor clustering by binding to both integrins and GFRs (***Hemler, 2005; Mikuličić et al., 2019***). In addition, earlier studies suggested that cofactors like laminin-332 and growth factors further stabilize these interactions (***Culp et al., 2006; Mikuličić et al., 2019; Richards et al., 2014; Surviladze et al., 2012***). Furthermore, the HPV capsid has multiple binding sites for distinct functions (***Richards et al., 2013***) which supports the idea that HS fulfills several roles during viral entry — first by promoting capsid attachment, then inducing conformational changes, and finally by assisting in receptor engagement and clustering. However, HS cleavage alone is not sufficient for ECM release; active transport and capsid priming remain essential (see leupeptin effect in Figure 2B and D).

### PsV-CD151 association occurs early at the cell surface

CD151 is a known mediator of HPV entry (***Scheffer et al., 2013; Spoden et al., 2008***). Our data shows that about 10% of the PsVs are closely associated with CD151 at all time points. This fraction is lower at 0 min post-CytD (Figure 5D, see also lower PCC in Figure 5C), indicating that actin disruption impairs initial CD151 engagement. Hence, the data suggest that PsVs establish contact to CD151 assemblies already during recruitment to the cell surface, and from then on, the fraction of closely associated PsVs remains constant. At first glance this implies that, after the initial formation of the PsV-CD151 assemblies, they do not change in nature. However, we observe agglomerated CD151 maxima (locally patched maxima) close to PsVs at later time points that likely are endocytic structures (see arrow in Figure 5A and Supplementary Figure 4). This agglomeration process is not detected in our analysis as we only determine the next nearest CD151 maximum. Some further CD151 reorganization into larger entry platforms has been previously observed (***Florin and Lang, 2018***). This is in accordance with the observation that virions enter the cell in crowds with many of them occupying one endocytic organelle (***Schelhaas et al., 2012***).

For the PsVs closely associated with CD151, one may have expected a larger fraction than 10%. Given that PsVs are not supposed to bind directly to CD151 and that CD151 is likely part of a larger membrane structure, like a tetraspanin enriched microdomain, our ≤ 80 nm distance criteria may be too short and by this we underestimate the associated fraction. Altogether, we propose that an early contact between PsVs and the cell surface involves CD151 (Figure 9).

**Figure 9.**
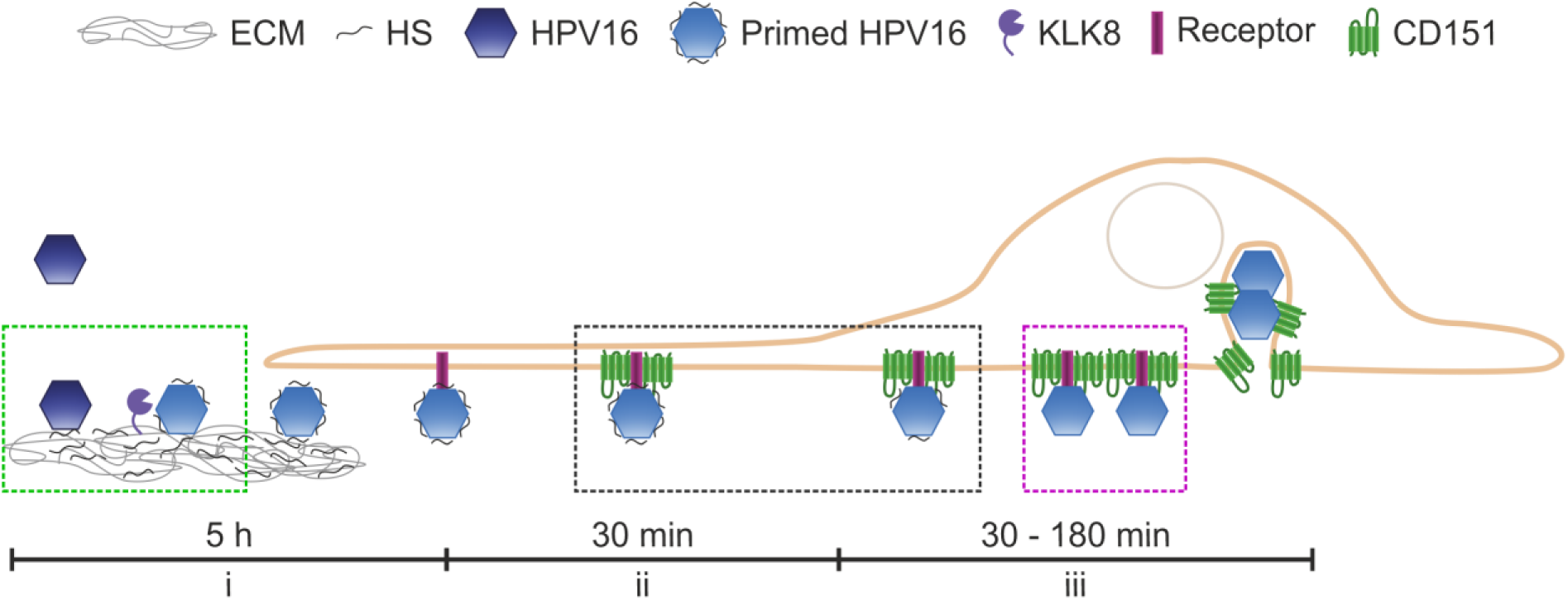
Model of ECM events, recruitment, and receptor engagement in HPV16 entry. (i) During 5 h of incubation with CytD, the PsVs bind to HS of the ECM, are primed, and become coated with HS cleavage products, enabling them for cell surface receptor engagement. After CytD removal, within 15 min, HS-decorated viruses are actively recruited to the cell body and (ii) associate with CD151 assemblies (completed within 30 min). (iii) Eventually, they lose their HS coat, and individual HPV16-CD151 assemblies agglomerate into larger structures (platforms), which are subsequently endocytosed. Dashed rectangles mark PsVs belonging to populations as defined in Figure 7. Dashed green rectangle, PsVs with a distance to HS < 250 nm and to Itgα6 > 250 nm. Dashed black rectangle, PsVs with a distance to HS < 250 nm and to Itgα6 < 250 nm. Dashed magenta rectangle, PsVs with a distance to HS > 250 nm and to Itgα6 < 250 nm.

### Integration of our data into the HPV infection cascade

HPV infection is the result of several steps, starting with the initial binding of virions via electrostatic and polar interactions (***Dasgupta et al., 2011***) to the primary attachment site HS (***Richards et al., 2013***), which induces capsid modification (***Cerqueira et al., 2015; Feng et al., 2024***) and HS cleavage (***Surviladze et al., 2015***), enabling the virion to be released from the ECM or the glycocalyx. Next, virions bind to the cell surface to a secondary receptor complex that forms over time, and become internalized via endocytosis, before they are trafficked to the nucleus (***Mikuličić et al., 2021; Ozbun and Campos, 2021***). Regarding the transition from the primary attachment site to cell surface receptor binding, as already outlined in the introduction, two models are discussed. In one model, proteases cleave the capsid proteins. After priming, the capsids are structurally modified and the virion can dissociate from its HS attachment site. It has been suggested that capsid priming is mediated by KLK8 (***Cerqueira et al., 2015***) and furin (***Richards et al., 2006***). In our system, KLK8 inhibition blocks PsV transport, while furin inhibition has some effect that, however, cannot be detected in this analysis (Figure 2D) suggesting furin engagement at later steps in the infection cascade. This is in line with earlier *in vitro* studies on the role of cell surface furin (***Day et al., 2008; Day and Schiller, 2009; Surviladze et al., 2015***). In any case, our results align with both models of ECM detachment: one involving HS cleavage (HS co-transfer) and another one involving capsid modification (by e.g., KLK8).

We propose that after 5 h of CytD treatment, glycan-induced structural activation as well as capsid processing by proteases such as KLK8 of the capsid and HS cleavage essentially are completed ((i) in Figure 9). Subsequently, the HS decorated virion is recruited from the ECM to the basal cell membrane, which takes about 15 min. In a time window of 30 min after CytD wash off (ii), PsV-CD151 association occurs. (iii) In the samples recorded 30 – 180 min after CytD wash off, we observe that PsVs lose their HS-coat, and individual PsV-CD151 assemblies seem to merge into larger structures that are subsequently endocytosed. This model highlights the role of ECM interactions, actin dynamics, and receptor engagement in HPV16 entry.

## Materials and Methods

### Antibodies and PsVs

We used the following primary antibodies in immunostainings of proteins. For the capsid protein L1, a rabbit polyclonal antibody (pAb) K75 (diluted 1:1000) as described previously (***Knappe et al., 2007; Rommel et al., 2005***), for CD151 a mouse monoclonal antibody (mAb) (1:200) (Biorad, cat# MCA1856GA), and for Itgα6 a rabbit polyclonal antibody (1:200) (Invitrogen, cat# PA5-12334) were used. For Heparan sulfate (HS) a mouse IgM monoclonal antibody (1:200) (amsbio, cat# 370255-S) was used that reacts with an epitope in native heparan sulfate chains and not with hyaluronate, chondroitin or DNA, and poorly with heparin (mAb 10E4 (***David et al., 1992***)). For HS neo-epitope (***Yokoyama et al., 1999***) detection, a mouse monoclonal antibody (1:200) (amsbio, cat#370260-S) was used that reacts only with heparitinase-treated heparan sulfate chains, proteoglycans, or tissue sections, and not with heparinase treated HSPGs. The antibody recognizes desaturated uronic acid residues (mAb 3G10 (***David et al., 1992***)). As secondary antibodies we used a STAR GREEN-coupled goat-anti-rabbit (1:200) (Abberior, cat# STGREEN-1002-500UG), an AlexaFluor 594-coupled donkey-anti-rabbit (1:200) (Invitrogen, cat# A21207), an AlexaFluor 594-coupled donkey-anti-mouse (1:200) (Invitrogen, cat# A21203), an AlexaFluor 594-coupled donkey-anti-mouse IgM (1:200) (Invitrogen, cat# A21044), a STAR RED-coupled goat-anti-mouse (1:200) (Abberior, cat# STRED-1001-500UG) and a STAR RED-coupled goat-anti-rabbit (1:200) (Abberior, cat# STRED-1002-500UG). Additionally, phalloidin iFluor488 (1:1000 of ready to use solution) (abcam, cat# ab176753) or phalloidin iFluor647 (1:1000) (abcam, cat# ab176759) was used to stain F-actin. Staining of the PsV plasmid DNA was performed by click-labeling according to the manufacturer’s instructions with the dye 6-FAM Azide (Baseclick EdU 488 kit, Carl Roth, cat# 1Y67.1).

HPV16 PsVs were produced following established procedures (***Buck et al., 2004; Spoden et al., 2012***). HEK293TT cells were cultured in 175 cm² flasks and transfected with polyethylenimine (PEI), using equimolar amounts of codon-optimized HPV16 L1/L2 (pShell-16L1wt-16L2wt) and a reporter plasmid (pGL4.20-puro-HPV16 LCR (***Wüstenhagen et al., 2018***)). For the production of 5-ethynyl-2’-deoxyuridine (EdU)-labeled PsVs, the culture medium was replaced 5 hours post-transfection with fresh medium containing 30 μM EdU. After 48 hours, cells were harvested, centrifuged, and washed twice with phosphate-buffered saline (PBS) supplemented with 9.5 mM MgCl₂ (1x PBS/ MgCl2). After the final centrifugation, the pellet was resuspended in 1x PBS/ MgCl2 containing 0.5% Brij58 (Sigma-Aldrich), and 250 units Benzonase (Merck Millipore), and incubated at 37°C for 24 hours on a rotating platform. Subsequently, lysates were chilled on ice and the NaCl concentration was adjusted to 0.85 M. After clarification by centrifugation, the supernatant was loaded onto an iodixanol (Optiprep) gradient consisting of 39%, 33%, and 27% layers (bottom to top). Gradients were equilibrated for 90 minutes at room temperature before ultracentrifugation at 55,000 rpm for 3.5 hours at 16°C. Fifteen fractions of 300 µl each were collected from the top and analyzed via luciferase reporter assay to identify peak fractions. PsV titers were determined based on packaged genomes (viral genome equivalents, vge) as previously described (***Spoden et al., 2012***). The concentrations of stock solutions were 7.7 x 10^6^ vge/µl (for microscopy experiments) and 14.1 x 10^6^ vge/µl (for the luciferase assay).

### Cell culture

For microscopy, human immortalized keratinocytes (HaCaTs) (Cell Lines Services, Eppelheim, Germany) were cultured in high glucose Dulbecco’s modified Eagle’s medium (DMEM + GlutaMAX, Gibco, cat# 61965-026) supplemented with 10% FBS (PAN Biotech, cat# P30-3031) and 1% Penicillin/Streptomycin (10,000 U/ml Penicillin, 10 mg/ml Streptomycin; PAN Biotech, cat# P06-07100) at 37 °C with 5% CO2. For the luciferase infection assay, cells were grown at 37°C and 5% CO2 in Dulbecco’s modified Eagle’s medium (DMEM + GlutaMAX, Thermo Fisher Scientific), supplemented with 10% FBS (Sigma-Aldrich), 1% minimum essential medium non-essential amino acids (MEM non-essential amino acids (Thermo Fisher Scientific)), and 5 µg/ml ciprofloxacin (Fresenius Kabi). For experiments, the antibiotics in the medium were omitted.

### Sample preparation and immunostaining

About 150,000 HaCaT cells were plated onto 25 mm diameter poly-L-lysine (PLL) coated (100 µg/ml PLL for 30 min) glass coverslips in 6-well plates and incubated for 24 h at 37°C and 5% CO2. The next day, cells were incubated at 37°C and 5% CO2 for 5 h with PsVs (46 vge/plated cell) and 10 µg/ml cytochalasin D (CytD; stock solution 10 mg/ml in dimethyl sulfoxide (DMSO)) (Life Technologies, cat# PHZ1063) or 30 µM (−)-blebbistatin (stock solution 13.68 mM in DMSO) (Sigma-Aldrich, cat# B0560-1MG) in DMEM supplemented with 10% FBS. For controls, we added the same amount of DMSO without CytD or blebbistatin. Cells were washed with PBS (137 mM NaCl, 2.7 mM KCl, 1.76 mM KH2PO4, 10 mM Na2HPO4, pH 7.4) and fresh DMEM supplemented with 10% FBS was added. Cells were incubated further at 37°C and 5% CO2 for 0 min (here the medium was added followed by immediate removal), 15 min, 30 min, 60 min and 180 min.

In another set of experiments, about 150,000 HaCaT cells were plated onto 25 mm diameter PLL coated glass coverslips in 6-well plates and incubated for 24 h at 37°C and 5% CO2. The next day, cells were incubated at 37°C and 5% CO2 for 5 h with PsVs (46 vge/plated cell) and 10 µg/ml CytD either together with 100 µM leupeptin (stock solution 100 mM in ddH2O) (Carl Roth, cat# CN33.1) or 5 µM Furin inhibitor I (stock solution 5 mM in DMSO) (Sigma-Aldrich, cat# 344930-1MG) in DMEM supplemented with 10% FBS. For controls, we used only 10 µg/ml CytD. Cells were washed with PBS and fresh DMEM supplemented with 10% FBS was added. Cells were incubated further at 37°C and 5% CO2 for 0 min, 30 min and 60 min.

To analyze PsV binding to detached HaCaT cells, about 350,000 HaCaT cells per well were plated in 6-well plates and incubated for 24 h at 37°C and 5% CO2. The next day, cells were detached by a 15 min incubation with 10 mM EDTA (in PBS, pH 7.4) at 37°C and 5% CO2. Detached cells were collected and incubated with 46 vge/cell in DMEM supplemented with 10% FBS under constant rotation at 4 °C for 1 h. Cells were washed three times with PBS at 4 °C to remove any unbound PsVs. Then, cells were seeded onto 25 mm diameter PLL coated glass coverslips in 6-well plates and incubated for 1 h at 37°C and 5% CO2.

Before staining, cells were washed twice with PBS and fixed at room temperature (RT) with 4% PFA in PBS for 30 min, unless membrane sheets were generated. In this case, the coverslips were placed in ice cold sonication buffer (120 mM KGlu, 20 mM KAc, 10 mM EGTA, 20 mM HEPES, pH 7.2) and a 100 ms ultrasound pulse at 100% power was applied. This was repeated until in total about 10 pulses were applied at different locations of the coverslip. Then, membrane sheets were fixed like cells at RT with 4% PFA in PBS for 30 min. PFA was removed, and residual PFA was quenched by 50 mM NH4Cl in PBS for 30 min. Afterwards, samples were blocked with 3% BSA in PBS for 30 min. Staining of PsVs was performed by click-labeling with the dye 6-FAM for 30 min at RT according to the manufacturer’s instructions. Then, samples were washed three times with PBS. In case of no PsV labeling by click-chemistry, directly after blocking, the respective primary antibodies were added: mouse IgM mAb against HS (1:200), mouse mAb against Δ-HS (1:200), rabbit pAb against Itgα6 (1:200), mouse mAb against CD151 (1:200) and rabbit pAb K75 against L1 (1:1000) in 3% BSA in PBS for 2 h. Samples were washed three times with PBS before adding the respective secondary antibodies and fluorescent labelled phalloidins in 3% BSA in PBS for 1 h: for HS we added AlexaFluor 594 coupled to donkey-anti-mouse IgM (1:200), for Δ-HS AlexaFluor 594 coupled to donkey-anti-mouse (1:200), for Itgα6 STAR RED coupled to goat-anti-rabbit (1:200), for CD151 AlexaFluor 594 coupled to donkey-anti-mouse (1:200) or STAR RED coupled to goat-anti-mouse (1:200), for L1 AlexaFluor 594 coupled to donkey-anti-rabbit (1:200) or STAR GREEN coupled to goat-anti-rabbit (1:200) and for F-actin phalloidin iFluor488 (1:1000) or phalloidin iFluor647 (1:1000). Afterwards, samples were washed three times with PBS, followed by mounting of the coverslips onto microscopy slides using ProLong Gold antifade mounting medium (Invitrogen, cat# P36930).

### Confocal and STED microscopy

For confocal and STED microscopy, samples were imaged employing a 4-channel STED microscope from Abberior Instruments (available at the superresolution light microscopy facility of the LIMES institute, Bonn, Germany). The microscope is based on an Olympus IX83 confocal microscope equipped with an UPlanSApo 100x (1.4 NA) objective (Olympus, Tokyo, Japan). Confocal and STED micrographs were recorded simultaneously. For confocal imaging, a 488 nm laser was used for the excitation of 6-FAM, STAR GREEN and phalloidin iFluor488, and emission was detected at 500–550 nm. For STED imaging, a 561 nm laser (at 45%) was used for the excitation of AlexaFluor 594 (detection at 580–630 nm) and a 640 nm laser (at 45%) for the excitation of STAR RED and iFluor647-labelled phalloidin (detection at 650-720 nm), in combination with a 775 nm laser for depletion (at 45%). The pixel size was set to 25 nm, for overview images a pixel size of 250 nm was used.

### Analysis of confocal and STED micrographs

Images were analyzed using Fiji ImageJ (***Schindelin et al., 2012***) in combination with a custom written macro, essentially as described previously (***Schmidt et al., 2024***). When analyzing double stainings of Alexa594 (red channel) and STAR RED (long red channel), we corrected for crosstalk from the red into the long red channel by subtracting 40% of the intensity of the red channel image from the long red channel image.

First, images were smoothed with a Gaussian blur to improve maxima detection. For confocal PsV-DNA micrographs, we employed a Gaussian blur of σ = 1, and for confocal PsV-L1 and STED micrographs a Gaussian blur of σ = 0.5. Local maxima were detected using the ‘Find Maxima’ function (noise tolerance 60 for L1 if stained together with phalloidin iFluor488, and 80 if stained together with CD151, HS, or phalloidin iFluor647, 3 for click-labeled PsVs, 8 for CD151, 6 for HS and 15 for Itgα6), yielding maxima positions in pixel positions. For analysis, regions of interest (ROIs) were defined (for details see figure legends).

In order to define the cell border region (Supplementary Figure 3B), the ImageJ ‘Make Binary’ function was used on CD151 STED micrographs for generating a binary mask. When possible, the ‘Wand’ tool was used to further outline the area covered by the cell, otherwise the same was done manually using the original micrograph as a reference. From the outline in the binary mask, a ROI was created, that was then filled in white with the ’Flood Fill’ Tool. The ROI was shrunk to a 20-pixel broad strip, and cleared outside of the strip, what leaves a 20-pixel broad, closed ribbon marking the intracellular side of the cell border that, however, exhibits arbitrary edges produced in the process above. These edges were removed by manual adjustments with the ’Pencil’ tool and the clear function. Afterwards, the ROI defines the intracellular side of the cell border. The ROI was symmetrically broadened by 40 pixels (using the ‘Enlarge’ function). From the now at least 60 pixels broad ROI, approximately two thirds covered the intracellular and one third the extracellular side. In this ROI, using the ImageJ ‘Find Maxima’ function, PsV-L1 maxima were detected (at a noise tolerance of 80) with a custom written macro in ImageJ both in the cell border region ROI and in a ROI covering the entire micrograph (excluding a 2-pixel edge). From these values, the percentage of PsVs within the cell border region was calculated.

For measuring intensity over time, we measured in the ROI the mean gray value. For background correction, we subtracted the mean gray value measured in a ROI next to the cell.

For measuring maxima intensity, a ROI with a diameter of 125 nm (5 pixels) was placed onto the determined maxima positions (see above). Using these ROIs, the average mean gray value of each maximum was measured. The average background mean gray value was measured in a ROI placed next to the cell and subtracted from the average mean gray value of the maxima. For each membrane sheet, the average maxima intensity was calculated, followed by averaging of the values of the membrane sheets.

To measure the shortest distance, e.g., between PsV and CD151 maxima, as a quality control for the PsV maxima (CD151 maxima do not undergo the quality control step) we placed onto each maximum position a horizontal and a vertical linescan (31 pixels long × 3 pixels width). Afterwards, we fitted a Gaussian distribution to the intensity distribution of each maximum. Only if at least one of the fits exhibits a fit quality of R^2^ > 0.8 and if the Gaussian maximum located central to the intensity distribution the maximum was included in the further analysis. Moreover, using a 125 nm ROI at the PsV and CD151 maxima positions, the center of mass of fluorescence was determined, yielding the maxima positions in sub-pixels. Based on these positions we further calculated the shortest distance of a PsV maximum to the nearest CD151 maximum.

Using the shortest distances, we either calculated the fraction of closely associated PsVs (PsVs with a distance ≤ 80 nm to e.g., CD151) over time, or the average shortest distance over time. The fraction of closely associated PsVs was corrected for random background association that increases with the maxima density. The relationship between background association and maxima density was expressed by a linear equation obtained from the same images re-analyzed after flipping them horizontally and vertically (see e.g., Supplementary Figure 7).

The Pearson correlation coefficient (PCC) was calculated with a custom written macro in ImageJ. As a control, we performed the analysis on horizontally and vertically flipped ROI-defined images as well (see e.g., Supplementary Figure 5A). The cell body PCC between PsVs and HS is measured in smaller ROIs covering exclusively the cell body region (this is confirmed with the Itgα6 or F-actin image as reference).

Per condition and biological replicate, 14 - 15 images were analyzed and values were averaged. For images from one set of experiments, in the figure the same channels are shown at the same settings of brightness and contrast, if not stated otherwise.

### Epi-fluorescence microscopy and image analysis

For epifluorescence microscopy, we used microscopic equipment and settings as previously described (***Homsi et al., 2014***) except of the illumination system, which was replaced by a SPECTRA X Light engine; Lumencor, Beaverton, OR, USA). In brief, PFA-fixed cells were imaged in a chamber filled with 1 ml of PBS to which 50 µl of a saturated solution of TMA-DPH [1-(4-tri-methyl-ammonium-phenyl)-6-phenyl-1,3,5-hexatriene-p-toluenesulfonate (T204; Thermofisher)] in PBS was added, for visualizing the membranes in the blue channel. PsVs, HS and Itgα6 were imaged in the green, red and far-red channels, respectively.

For analysis, rectangular ROIs of ≈ 45 x 10^3^ pixel were placed onto the cell body such that approximately one half of the ROI covers the ECM area and the other one the cell body. The cell periphery was defined using the TMA-DPH image from which a binary mask was created. With reference to the binary mask, a band of up to 30 pixels width was created beginning at the cell body and reaching out towards the cell periphery. Using this ROI, in the PsV-DNA image the size of the ROI and the mean gray values were measured from which the integrated intensities were calculated after background correction.

### Luciferase infection assay

HaCaT cells were grown in 24-well plates and allowed to adhere overnight. The following day, cells were treated with HPV16 PsVs at a concentration of approximately 100 vge per cell, in the presence or absence of 10 µg/ml cytochalasin D (CytD) in DMSO or an equivalent amount of DMSO (Control). In one condition, PsVs/CytD were applied for 5 hours, after which the medium was replaced with fresh medium lacking the compounds, and incubation was continued for additional 19 hours (in total 24 h). In another condition, cells were exposed to PsVs/CytD continuously for the full 24 h period. Then, cells were washed with PBS and lysed using 1 x Cell Culture Lysis Reagent (Promega, cat# E153A). Following high-speed centrifugation, luciferase activity in the cleared lysates was quantified using an LB 942 Tristar 3 Multimode Microplate Reader (Berthold Technologies). Cytotoxic effects, accompanied by a loss of membrane integrity (indicated by released lactate dehydrogenase (LDH)), were determined by measuring the LDH levels in the cell lysates. LDH activity was assessed using the CytoTox-ONE Homogeneous Membrane Integrity Assay (Promega, cat# G7891). LDH fluorescence was quantified according to the manufacturer’s instructions using the LB 942 Tristar 3 Multimode Microplate Reader.

### Statistics

Microscopy data sets were based on three biological replicates. One replicate includes per time point the average of 14 - 15 images analyzed (with exception of Figure 4, which includes 14 – 35 analyzed cells per replicate and time point). For the infection assay, data sets were based on three biological replicates. One biological replicate is the average of three technical replicates. Data was tested for significance with a two-tailed, unpaired student’s *t*-test with significance * = p < 0.05, ** = p < 0.01 and *** = p < 0.001.

## Acknowledgement

LF and TL were funded by the Deutsche Forschungsgemeinschaft (DFG, German Research Foundation), Projektnummer 322863883 (LF and TL) and Projektnummer 270976260 (TL). This publication was supported by the Open Access Publication Fund of the University of Bonn.

## Supplementary information

**Supplementary Figure 1.**
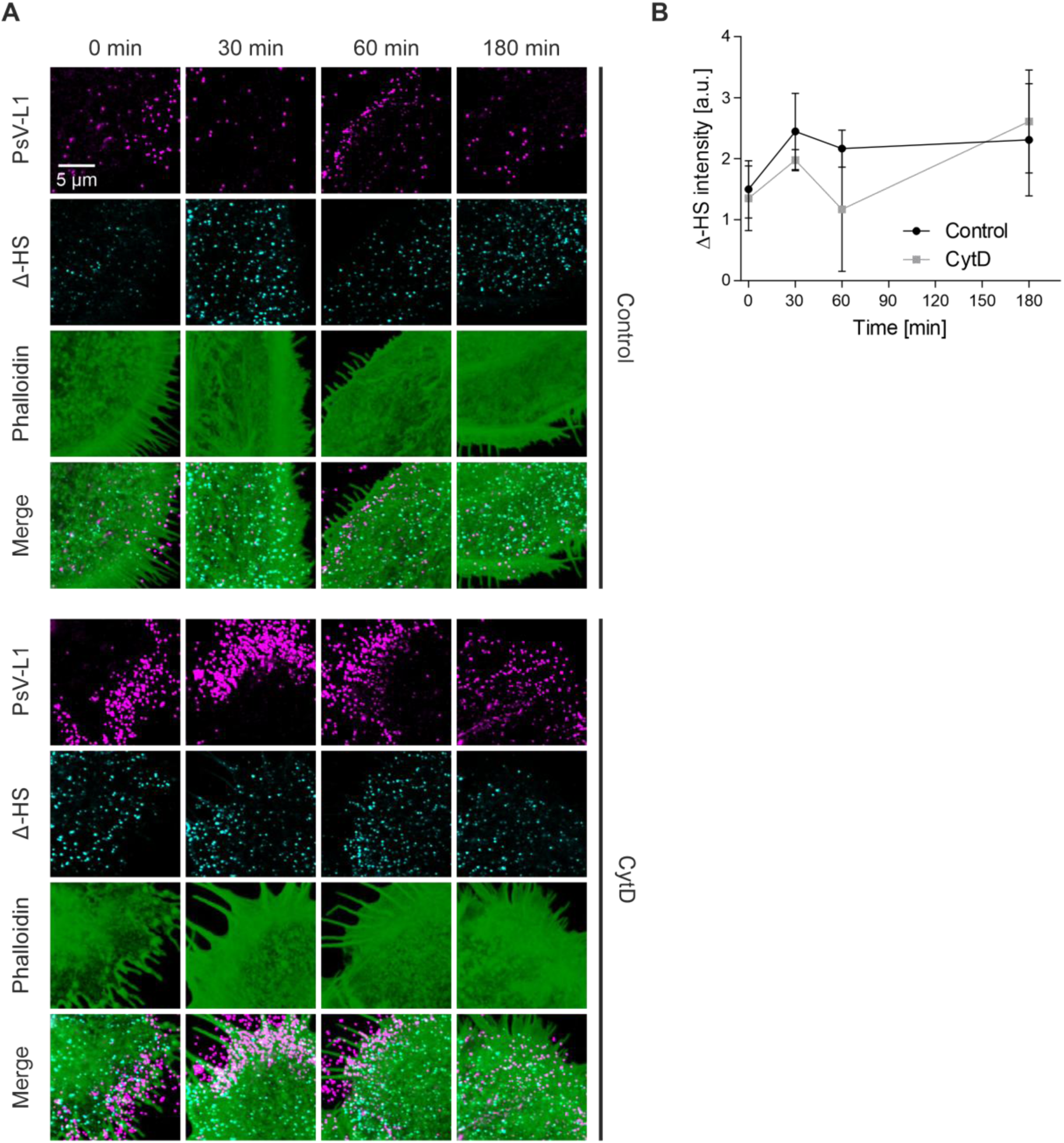
HS neo-epitope (Δ-HS) staining after CytD treatment. (A) HaCaT cells were incubated with PsVs at 37 °C for 5 h, in the absence (Control, upper panels) or presence of 10 µg/ml CytD (CytD, lower panels). Afterwards, cells were washed and incubated without PsVs/CytD further for up to 180 min, before they were fixed and stained. Immunofluorescence was used for the visualization of L1 (STAR GREEN; magenta LUT) and the HS neo-epitope, referred to as Δ-HS (AlexaFluor 594; cyan LUT). F-actin was stained by iFluor647-labelled phalloidin (green LUT). PsV-L1, Δ-HS and F-actin staining were imaged in the confocal mode of a STED microscope. For analysis, we placed large ROIs onto the images that covered mainly the cell body but included parts of the cell periphery as well. (B) The mean Δ-HS intensity over time. Values are given as means ± SD (n = 3 biological replicates). Statistical differences between the same time points of Control and CytD were analyzed by using the two-tailed, unpaired student’s *t*-test (n = 3), but the analysis yielded no p-values below 0.05.

**Supplementary Figure 2.**
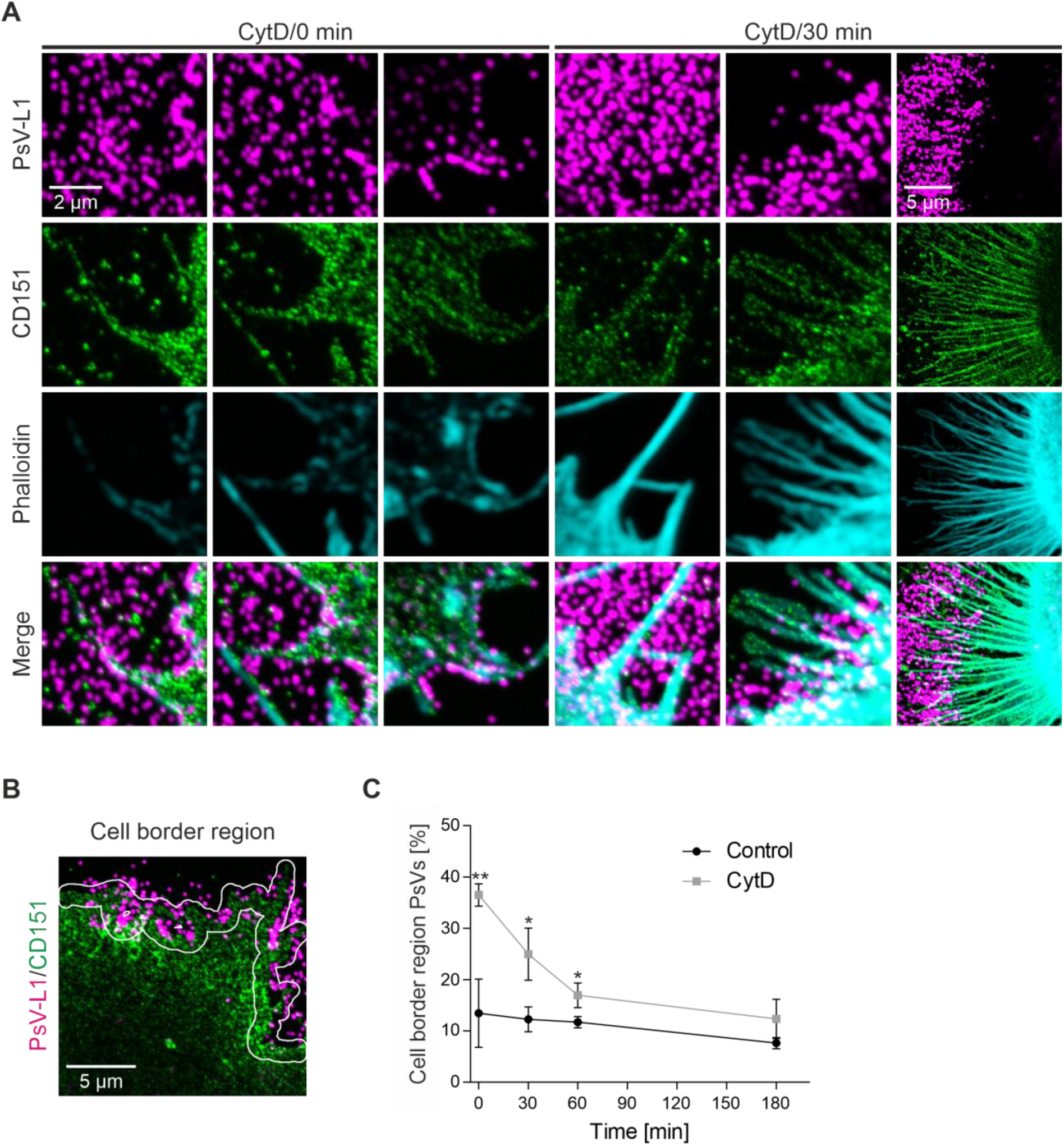
Variability of filopodia/diminishment of PsVs from the cell border region after CytD removal. (A) More images from the experiment described in Figure 5. We observed that many cells exhibit filopodia. Due to the large variability in number and shape we cannot show a representative image. Instead, we present for the CytD condition (0 min and 30 min) a variety of cells with CD151 positive filopodia. (B) Based on the CD151 image (green LUT), a cell border strip is defined and broadened by 20 pixels on each side, in the following referred to as the cell border region (for details see materials and methods). Please note that the cell border region is different from the cell periphery described in Figure 4 that delineates the cell body from the periphery. Instead, the cell border region covers both cell body and periphery (approximately two thirds inside and one third outside of the cell; for details see methods). (C) PsV maxima in the cell border region over time, expressed as percentage of all PsVs present in the image. Values are given as means ± SD (n = 3 biological replicates). Statistical differences between the same time points of Control and CytD were analyzed by using the two-tailed, unpaired student’s *t*-test (n = 3, for details see materials and methods).

**Supplementary Figure 3.**
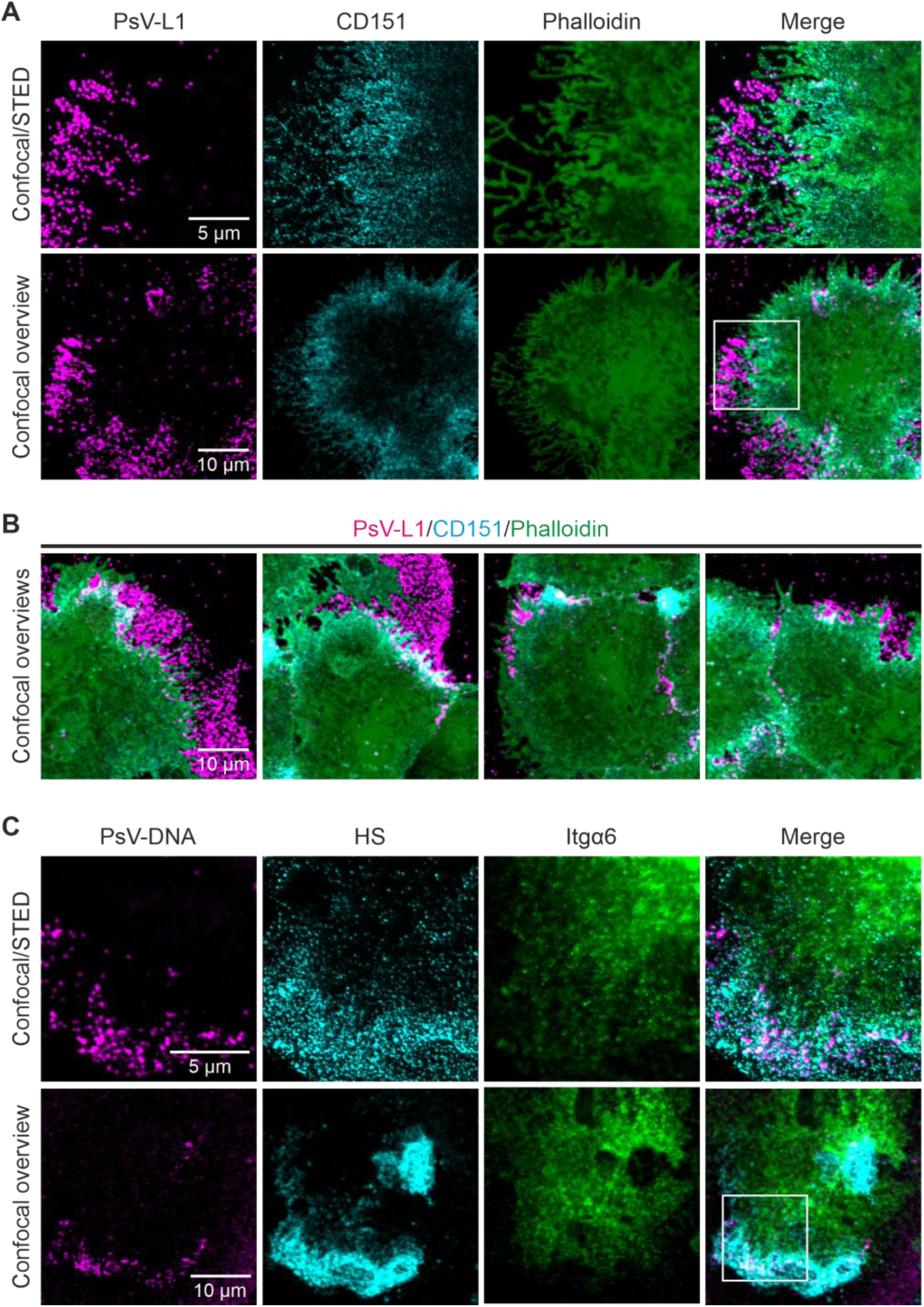
Overview images. (A) Upper row, micrographs shown in Figure 5A, CytD/0min, using for CD151 a cyan instead of a green LUT, and showing as well F-actin staining (green LUT). Lower row, confocal overview images. After taking the overview images, the area in the white box was imaged as described in the legend of Figure 5, yielding the images shown in the upper row. (B) More merged overview images from the experiment described in Figure 5, condition CytD/0 min, showing examples of large and small PsV accumulations. (C) Upper row, same micrographs as shown in Figure 6A, CytD/0 min. Lower row, confocal overview images. The area in the white box was imaged as described in the legend of Figure 6, yielding the images of the upper row in C.

**Supplementary Figure 4.**
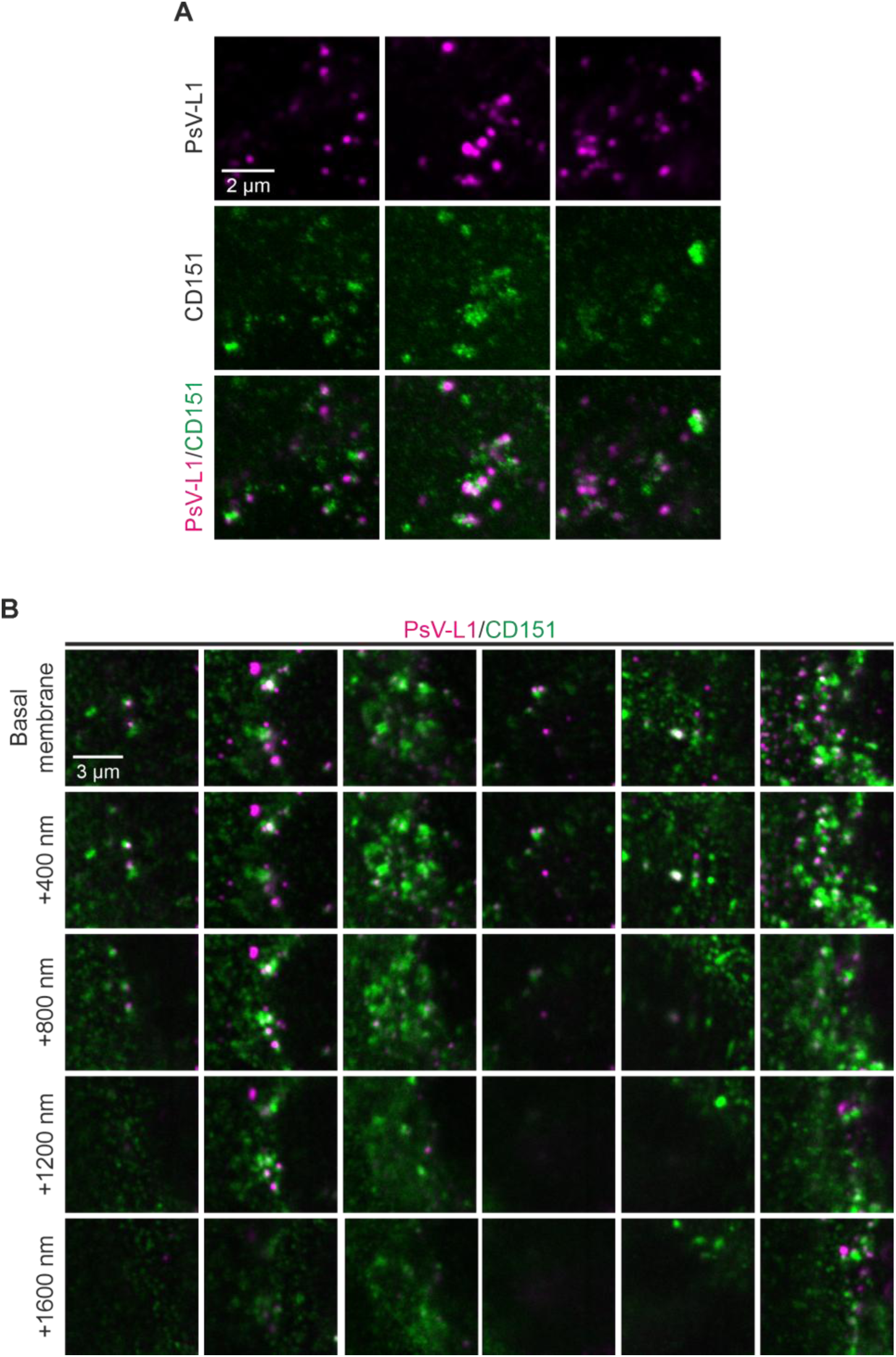
Examples of agglomerated CD151 maxima associated with PsVs that presumably represent endocytic structures. (A) From the CytD/180 min time point described in Figure 5, we show more examples of agglomerated CD151 maxima (green LUT, see green patches) that associate with PsVs (magenta LUT). (B) Same experiment as in (A). Starting at the basal membrane, at confocal resolution we imaged CD151 (green LUT) and PsV-L1 (magenta LUT). Cells were further scanned by 400 nm steps in the axial direction. Some of the CD151 agglomerates noticed in the basal membranes appear to continue deeper into the cell, in the second example from the left more than a micrometer. We propose that the agglomerated CD151 maxima that overlap with PsVs are endocytic structures, as CD151 has been shown to co-internalize with PsVs (***Scheffer et al., 2013***), and as these structures invaginate into the cell, like PsV filled tubular organelles previously described by electron microscopy (***Schelhaas et al., 2012***). Images in (A) and (B) are shown at the same settings of brightness and contrast but at different settings compared to Figure 5.

**Supplementary Figure 5.**
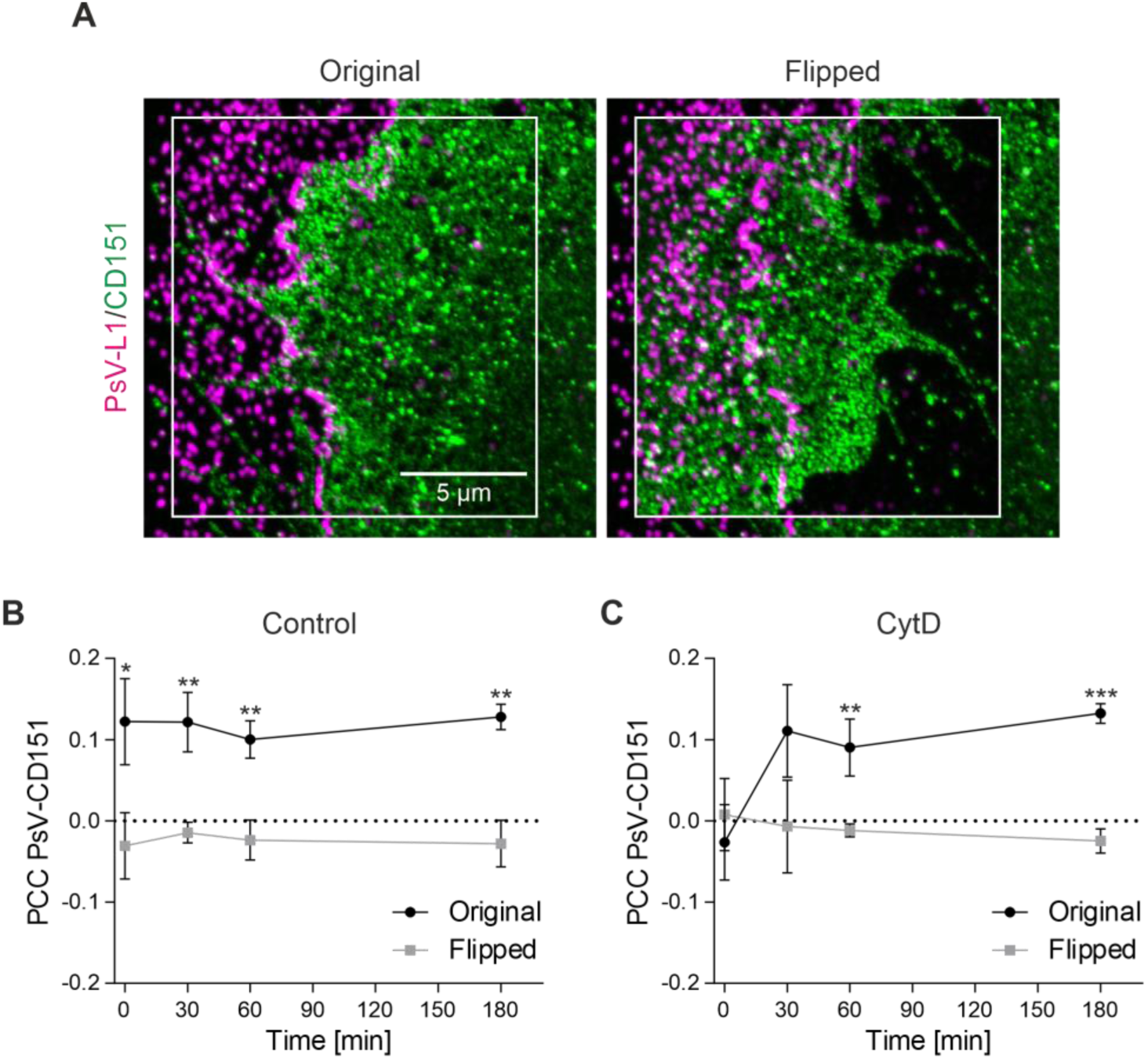
PCC values of flipped images. (A) For each image pair, we determined the PCC on non-overlapping images with the same density of objects and intensities. To this end, one channel was flipped horizontally and vertically. Left, original images (from the CytD/0 min condition). The ROI for analysis is illustrated as white box. Right, in the ROI, the CD151 image (green LUT) was flipped horizontally and vertically. (B + C) Shown are again the PCCs over time of Figure 5C of the original images ((B) Control, (C) CytD) together with the PCCs of the respective flipped images. Values are given as means ± SD (n = 3 biological replicates). Statistical differences between the same time points of original and flipped images was analyzed by using the two-tailed, unpaired student’s *t*-test (n = 3, for details see methods).

**Supplementary Figure 6.**
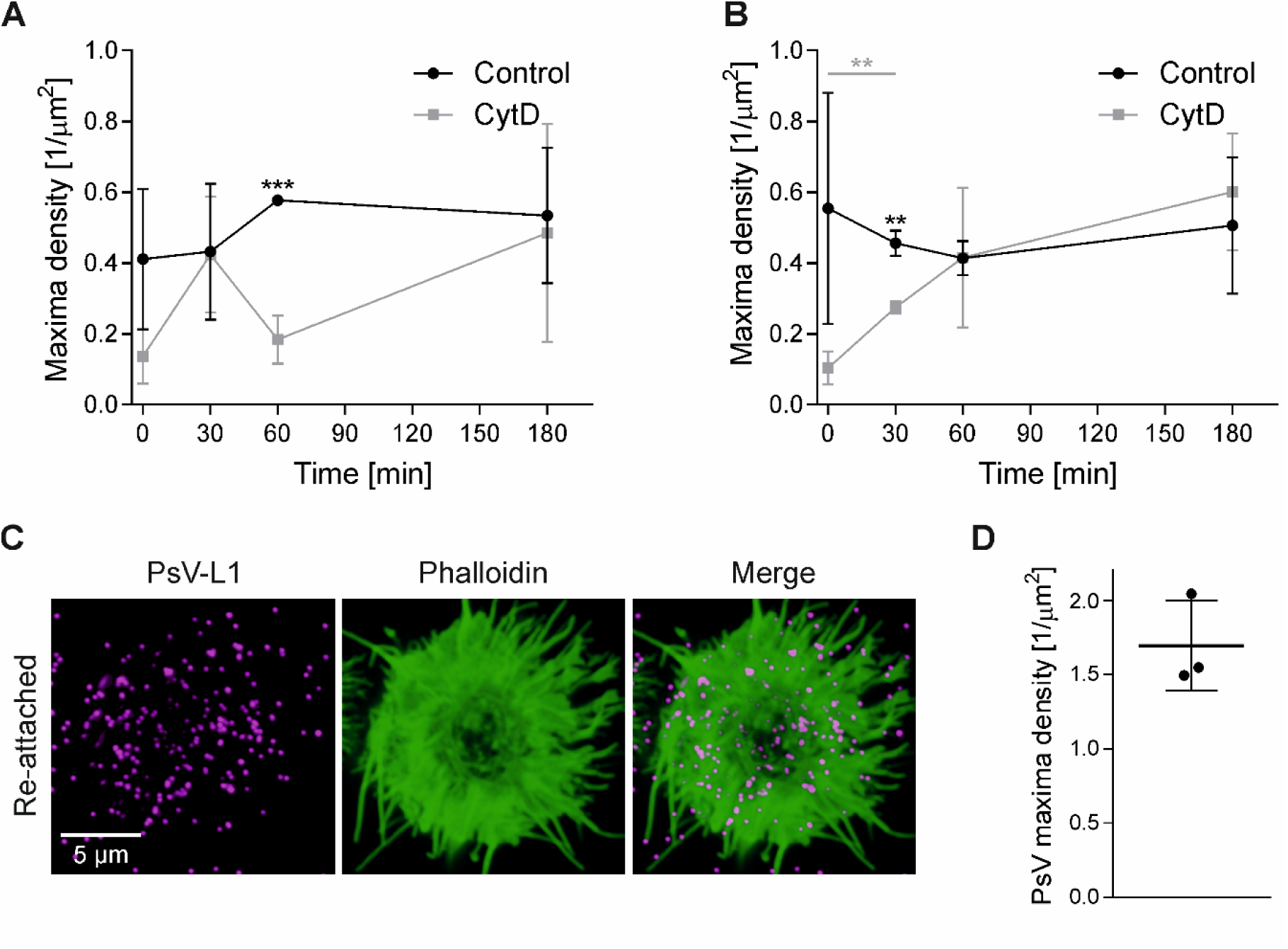
PsV density at the membrane of normally grown and re-attached HaCaT cells. (A + B) In order to count the PsVs at the basal membrane of the experiments shown in (A) Figure 5 (PsVs visualized by antibody labeling) and (B) Figure 6 (PsVs visualized by click-chemistry), we placed ROIs covering only the cell body. Within these ROIs the PsV maxima were counted and related to the size of the analyzed area. Values are given as means ± SD (n = 3 biological replicates). Statistical differences between Control and CytD at each time point were analyzed by using the two-tailed, unpaired student’s *t*-test (n = 3, for details see methods). (C) HaCaT cells were detached by a 15 min incubation with 10 mM EDTA (in PBS) and incubated with PsVs at 4 °C for 1 h under constant rotation. After washing the cells three times with PBS, they were seeded onto PLL-coated glass coverslips for 1 h. Then, cells were washed, fixed and stained for L1 (STAR GREEN; magenta LUT) by indirect immunofluorescence and for F-actin by iFluor647-labelled phalloidin (green LUT). PsV-L1 and F-actin staining were imaged as in Figure 5 in the confocal mode of a STED microscope. (D) Analysis as in (A + B). Values are given as means ± SD (n = 3 biological replicates).

**Supplementary Figure 7.**
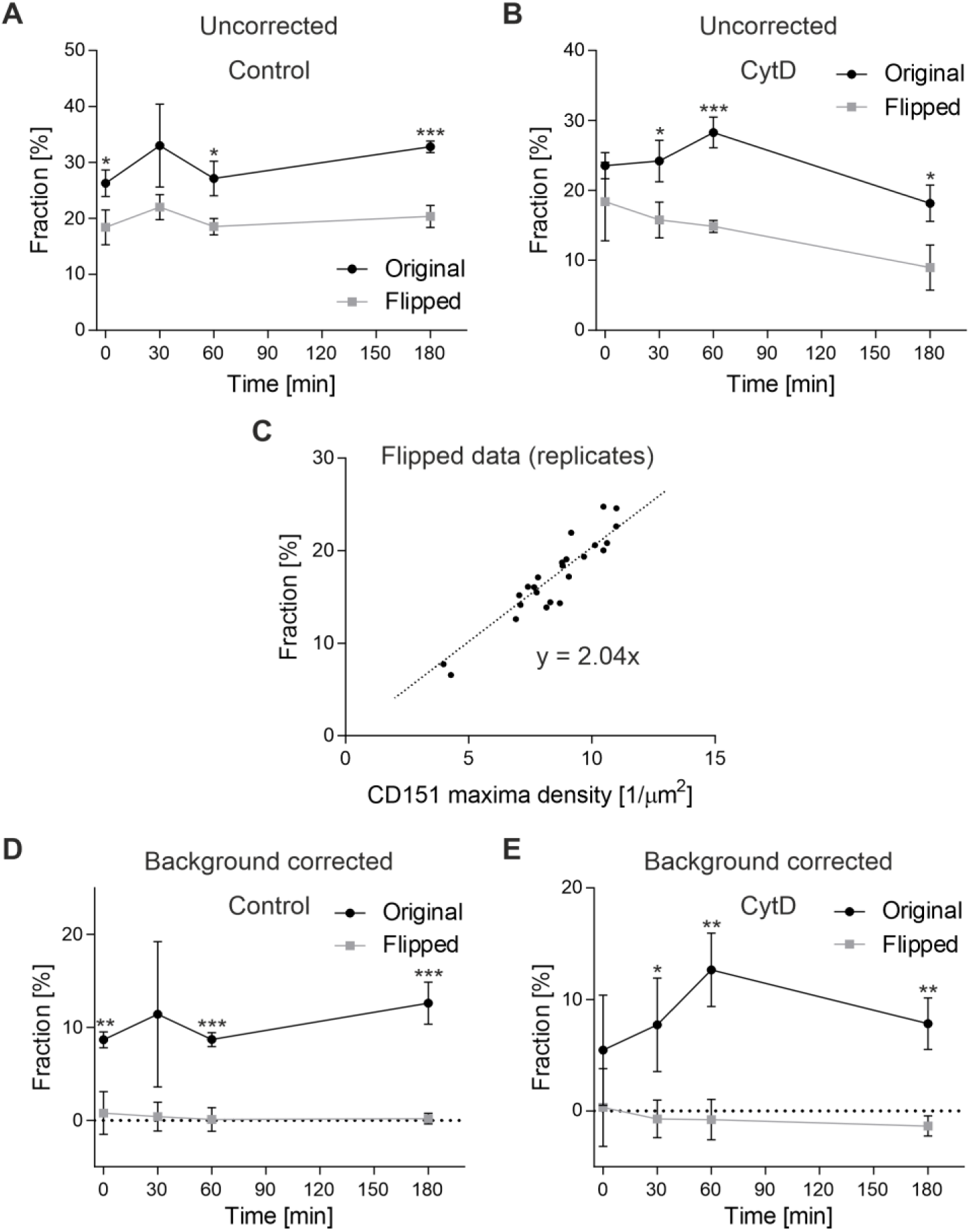
Background correction of the fraction of PsVs closely associated with CD151. (A) The fraction of closely associated PsVs (PsV-L1 maxima with a distance ≤ 80 nm to the next nearest CD151 maximum) in the Control of Figure 5 was analyzed on original and flipped images (for an example of a flipped image see Supplementary Figure 5A). As shown in (A), on flipped images, we often find values more than half of the values of the original images, demonstrating that many PsVs have a distance ≤ 80 nm to CD151 merely by chance, in the following referred to as background association. (B) Same as (A), for CytD. (C) The value of each time point in (A) and (B) is the average of three biological replicates. We take the altogether 24 fraction values obtained on flipped images (12 values from Control and CytD each), and plot the fraction of closely associated PsVs against the average CD151 maxima density in the respective images. As can be seen in (C), the fraction increases with the maxima density, as the chance of a distance ≤ 80 nm increases with the maxima density. The fitted linear regression line describes how the background association depends from the maxima density. As a result, the background association (y) can be calculated for any maxima density (x) with the equation y = 2.04 • x. The CytD/0 min condition may be overcorrected, if it includes many images with CD151 flipped onto peripheral PsVs that actually are distal to CD151 (for an example ROI see Supplementary Figure 5A). On the other hand, PsVs right at the cell border, where CD151 staining tends to be strong (Supplementary Figure 5A), after flipping have less CD151 than before, contributing to undercorrection. (D) and (E), for each original and flipped replicate included in (A) and (B), the background was calculated using the above equation and subtracted from the uncorrected value. The average original background corrected values are shown in Figure 5D. Values are given as means ± SD (n = 3 biological replicates). Statistical differences between the same time points of original and flipped images were analyzed by using the two-tailed, unpaired student’s *t*-test (n = 3, for details see methods).

**Supplementary Figure 8.**
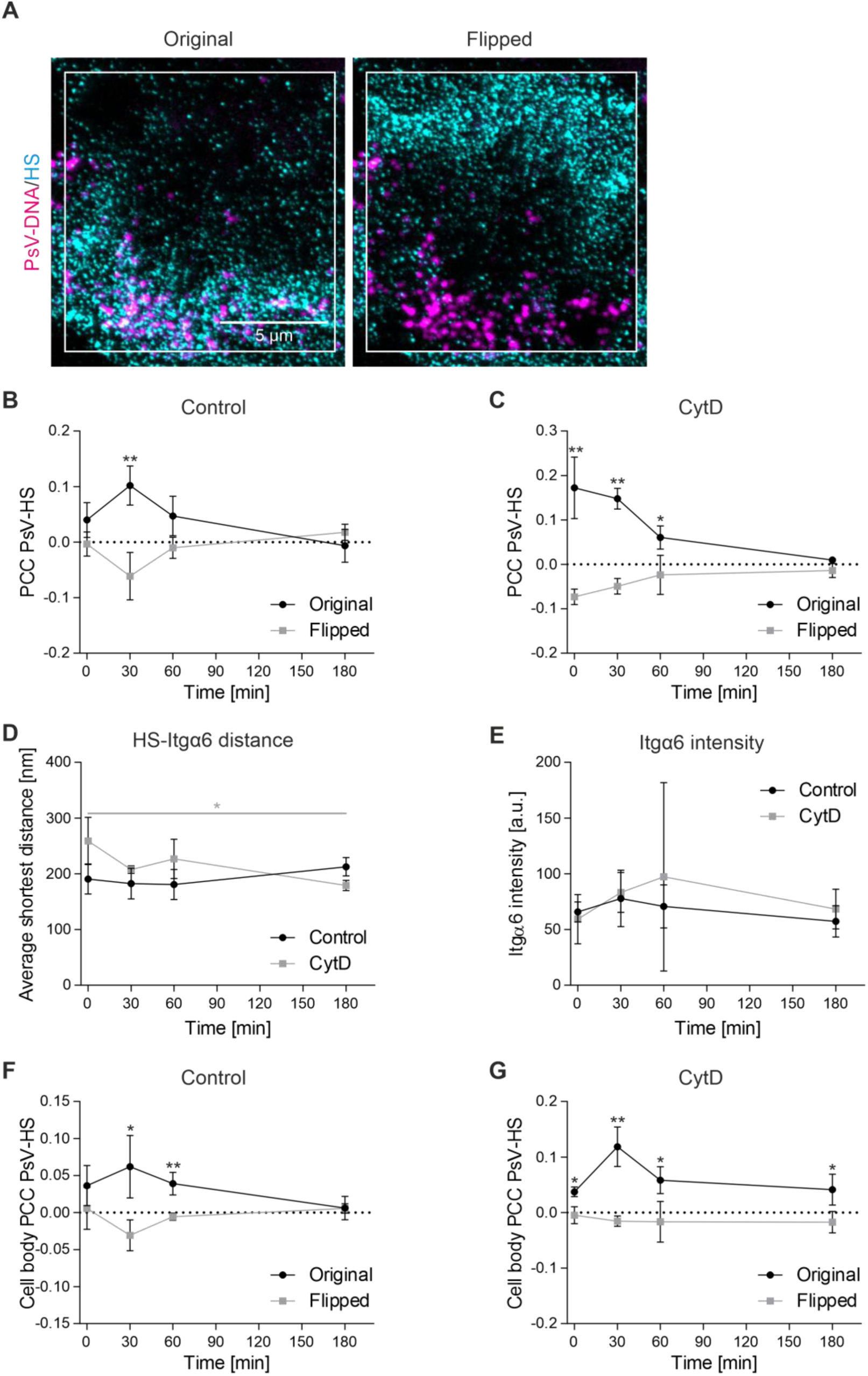
PCC values of flipped images, HS-Itgα6 shortest distance and Itgα6 intensity. (A) For each image pair, we determined the PCC on respective non-overlapping images with the same density of objects and intensities. To this end, one channel was flipped horizontally and vertically. Left, original images (from the CytD/0 min condition). The ROI for analysis is illustrated as white box. Right, in the ROI, the HS image (cyan LUT) was flipped horizontally and vertically. (B + C) Shown are the PCCs over time of Figure 6C obtained with the original images ((B) Control, (C) CytD) together with the PCCs of the flipped images. (D) The shortest distance of HS maxima to the next nearest Itgα6 maxima over time. (E) The mean Itgα6 intensity over time. (F + G) Shown are the PCCs over time of Figure 6D obtained with the original images ((F) Control, (G) CytD) together with the PCCs of the flipped images. Values are given as means ± SD (n = 3 biological replicates). Statistical differences between the same time points of original and flipped image ((B), (C), (E), (F) and (G)) or between CytD/0 min and CytD/180 min (D) were analyzed by using the two-tailed, unpaired student’s *t*-test (n = 3, for details see methods).

**Supplementary Figure 9.**
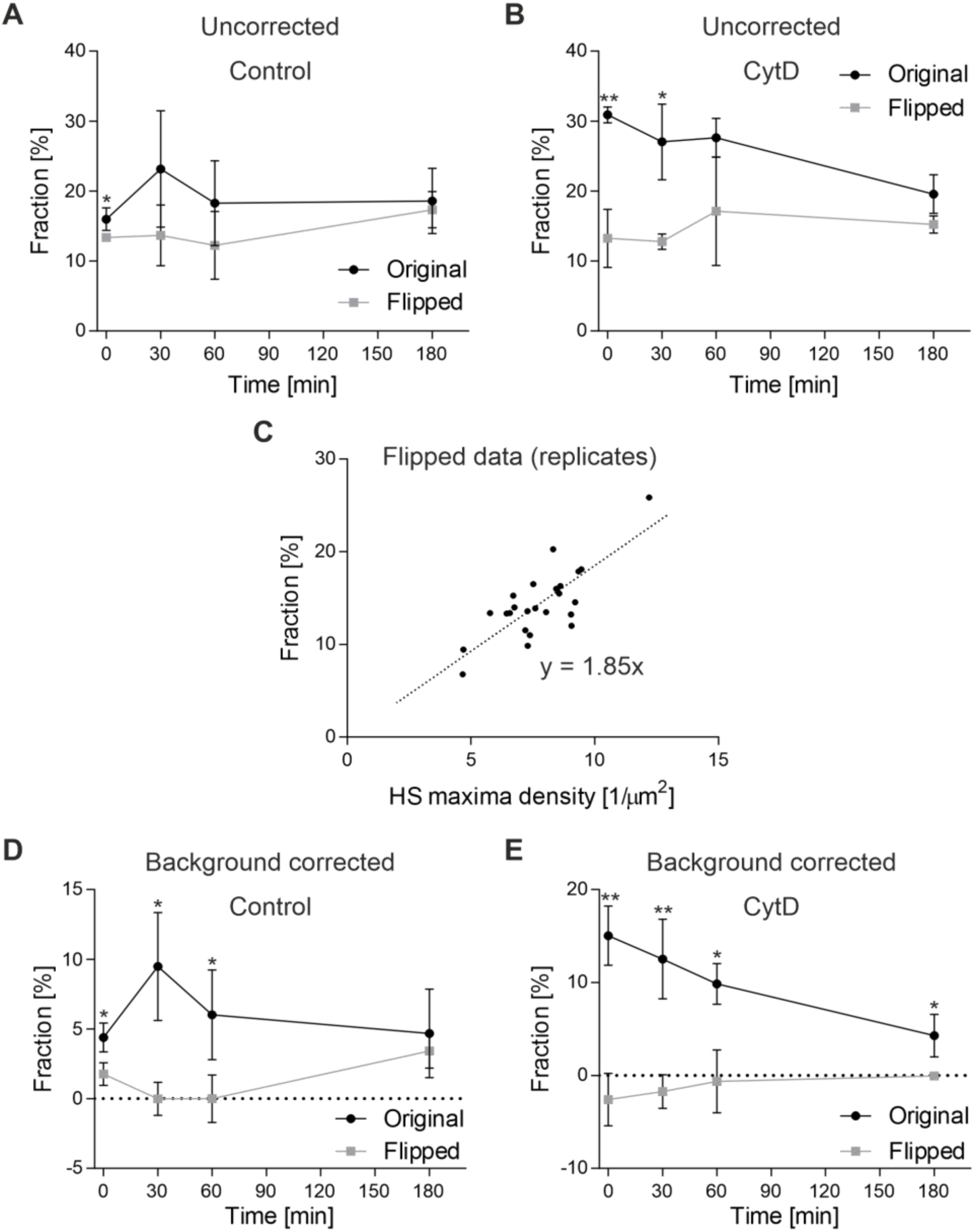
Background correction of the fraction of PsVs closely associated with HS. (A) The fraction of closely associated PsVs (PsV-DNA maxima with a distance ≤ 80 nm to the next nearest HS maximum) in the Control of Figure 6 was analyzed on original and flipped images (for an example of a flipped image see Supplementary Figure 8A). (B) Same as (A), for CytD. (C) From the altogether 24 fraction values obtained on flipped images (12 values from Control and CytD each), we obtain the equation y = 1.85 • x for calculating the background association (please see Supplementary Figure 7 for more details). (D) and (E), for each original and flipped replicate included in (A) and (B), the background was calculated using the above equation and subtracted from the uncorrected value. The average original background corrected values are shown in Figure 6E. Values are given as means ± SD (n = 3 biological replicates). Statistical differences between the same time points of original and flipped images were analyzed by using the two-tailed, unpaired student’s *t*-test (n = 3, for details see methods).

**Supplementary Figure 10.**
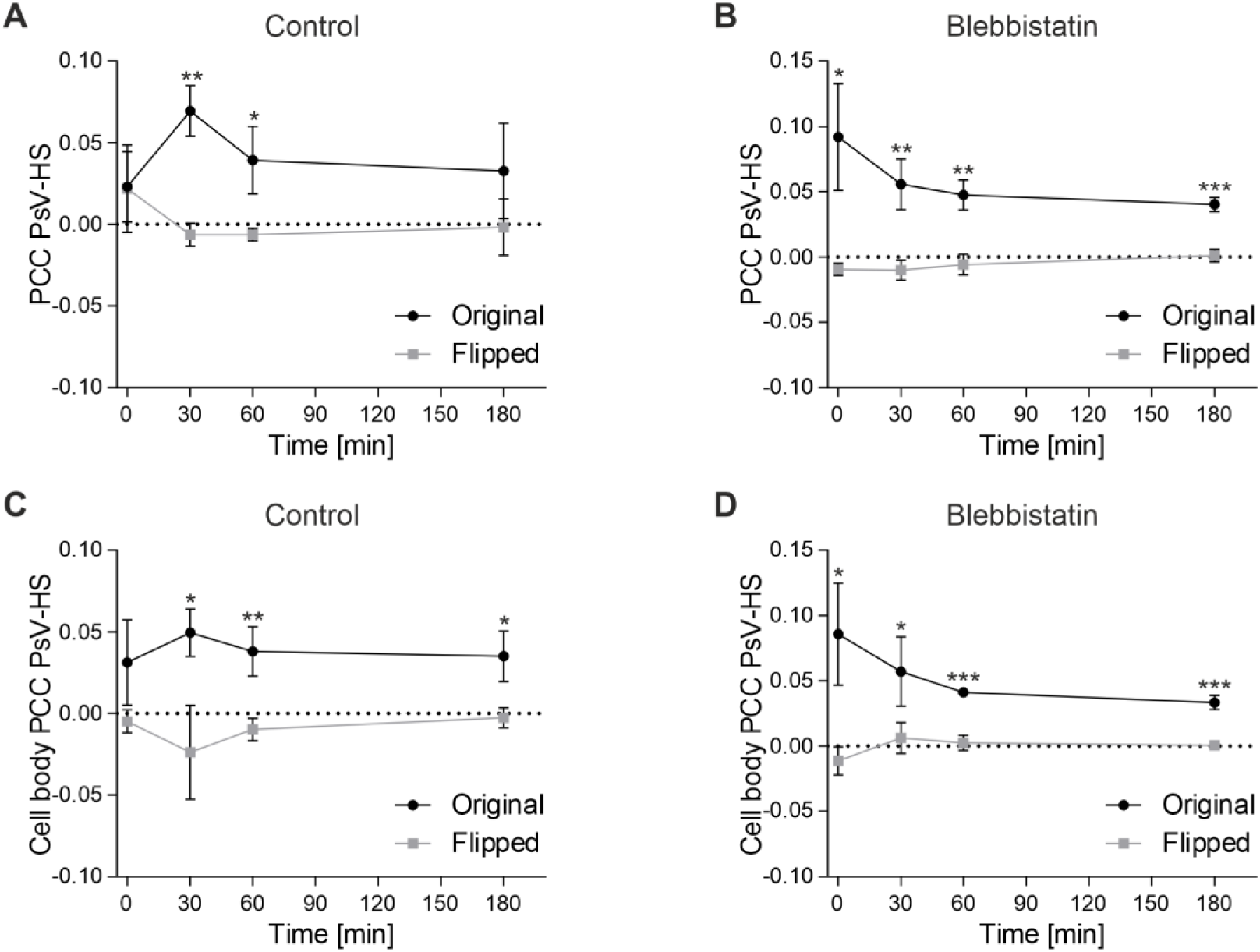
PCC values of flipped images. (A) and (B) The PCC values between PsV and HS of Figure 8C ((A) Control, (B) Blebbistatin) plotted together with the respective PCC values of flipped images. (C) and (D) The cell body PCC values between PsV and HS of Figure 8D ((C) Control, (D) Blebbistatin) plotted together with the respective PCC values of flipped images. Values are given as means ± SD (n = 3 biological replicates). Statistical differences between the same time points of original and flipped images were analyzed using the two-tailed, unpaired student’s *t*-test (n = 3, for details see methods).

**Supplementary Figure 11.**
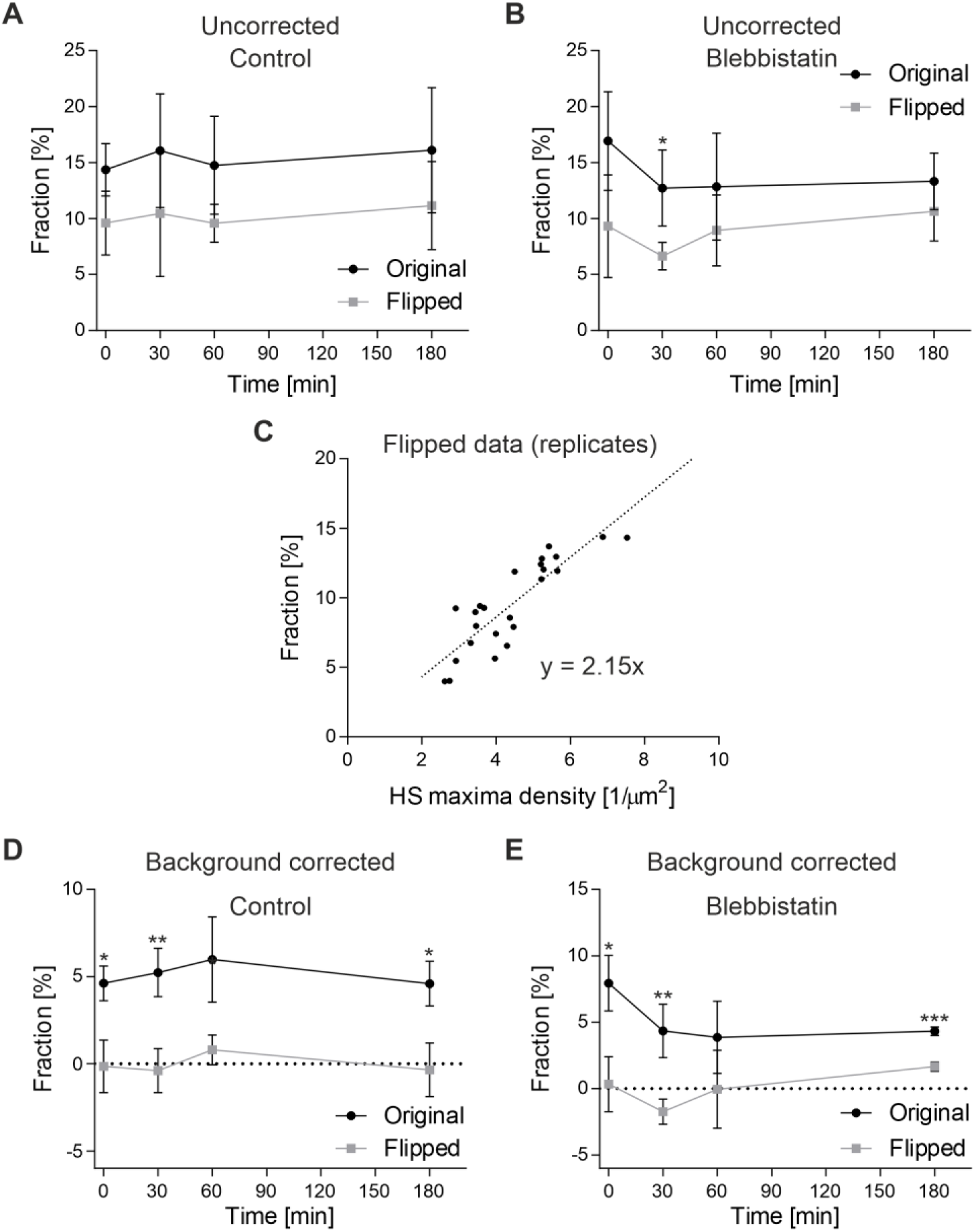
Background correction of the fraction of PsVs closely associated with HS after blebbistatin treatment. (A) The fraction of closely associated PsVs (PsV-L1 maxima with a distance ≤ 80 nm to the next nearest HS maximum) in the Control of Figure 8 was analyzed on original and flipped images. (B) Same as (A), for blebbistatin. (C) From the altogether 24 fraction values obtained on flipped images (12 values from Control and Blebbistatin each), we obtain the equation y = 2.15 • x for calculating the background association (please see Supplementary Figure 7 for more details). (D) and (E), for each original and flipped replicate included in (A) and (B), the background was calculated using the above equation and subtracted from the uncorrected value. The average original background corrected values are shown in Figure 8E. Values are given as means ± SD (n = 3 biological replicates). Statistical differences between the same time points of original and flipped images were analyzed by using the two-tailed, unpaired student’s *t*-test (n = 3, for details see methods).

**Supplementary Table 1.** Fraction of PsVs in percent of each of the four distance categories (see left column). Values are means ± SD of the data shown in Figure 7B. For each time point and category, p-values between Control and CytD were calculated by using the two-tailed, unpaired student’s *t*-test (n = 3 biological replicates). P-values < 0.05 are illustrated in bold.

|  | 0 min |  |  | 30 min |  |  | 60 min |  |  | 180 min |  |  |
| --- | --- | --- | --- | --- | --- | --- | --- | --- | --- | --- | --- | --- |
|  | Control [%] | CytD [%] | p-value | Control [%] | CytD [%] | p-value | Control [%] | CytD [%] | p-value | Control [%] | CytD [%] | p-value |
| HS < 250 nm & Itga6 < 250 nm | 62.31<br>± 5.05 | 47.83<br>± 3.26 | **0.0271** | 65.48<br>± 8.35 | 63.67<br>± 1.14 | 0.7764 | 59.06<br>± 10.64 | 67.16<br>± 9.84 | 0.4739 | 63.69<br>± 4.27 | 59.28<br>± 3.17 | 0.3065 |
| HS < 250 nm & Itga6 > 250 nm | 12.18<br>± 5.51 | 46.04<br>± 3.67 | **0.0019** | 15.46<br>± 3.14 | 26.12<br>± 2.16 | **0.0167** | 13.25<br>± 1.94 | 19.87<br>± 3.12 | 0.0637 | 11.97<br>± 1.53 | 10.64<br>± 1.15 | 0.3808 |
| HS > 250 nm & Itga6 < 250 nm | 21.59<br>± 4.07 | 4.34<br>± 0.37 | **0.0040** | 15.83<br>± 5.73 | 8.58<br>± 2.33 | 0.1723 | 23.49<br>± 6.55 | 9.60<br>± 4.81 | 0.0731 | 20.88<br>± 3.31 | 25.24<br>± 1.96 | 0.1842 |
| HS > 250 nm & Itga6 > 250 nm | 3.93<br>± 0.30 | 1.79<br>± 0.60 | **0.0110** | 3.23<br>± 0.88 | 1.63<br>± 0.21 | 0.0675 | 4.19<br>± 2.43 | 3.38<br>± 2.20 | 0.7425 | 3.46<br>± 0.54 | 4.84<br>± 2.87 | 0.5393 |

